# Platelet-derived transcription factors license human monocyte inflammation

**DOI:** 10.1101/2022.08.10.503291

**Authors:** Ibrahim Hawwari, Lukas Rossnagel, Nathalia Sofia Rosero Reyes, Salie Maasewerd, Marius Jentzsch, Agnieszka Demczuk, Lino L Teichmann, Lisa Meffert, Lucas S. Ribeiro, Sebastian Kallabis, Felix Meissner, Magali Noval Rivas, Moshe Arditi, Damien Bertheloot, Bernardo S. Franklin

## Abstract

CD14^+^ monocytes, the predominant population in human blood, are primarily engaged in host defense and pro-inflammatory cytokine responses. Aberrant monocyte activity causes life-threatening cytokine storms, while dysfunctional monocytes lead to ’immunoparalysis.’ Understanding the mechanisms controlling monocyte functions is therefore paramount. Here, we reveal platelets’ vital role in human monocytes’ pro-inflammatory responses. Low platelet counts in immune thrombocytopenia (ITP) patients, or platelet depletion in healthy monocytes result in monocyte immunoparalysis, characterized by reduced pro-inflammatory gene expression and weakened cytokine responses to immune challenge. Remarkably, adding fresh platelets reverses monocyte immunoparalysis. In mice, thrombocytopenia results in down-regulation of myeloid innate immune genes, and compromised host defense transcriptional programs in monocytes despite normal responses to LPS. Platelets control monocyte cytokines independently of traditional cross-talk pathways, acting as reservoirs of transcription factors like NFκB and MAPK p38. We pinpointed megakaryocyte-derived NFκB2 transfer to human monocytes by mass spectrometry-based proteomics. Functionally, platelets proportionally restored impaired cytokine secretion in human monocytes lacking p38a and NFκB. We unveil the intercellular transfer of inflammatory regulators, positioning platelets as central checkpoints in monocyte-mediated inflammation.

**Key Points:**

- Platelets are essential to TLR and NLR cytokine responses of human monocytes,
- Immune thrombocytopenia leads to monocyte immunoparalysis;
- Platelet supplementation reverses monocyte immunoparalysis;
- Platelets transfer NFκB that reactivates cytokine production in genetically deficient monocytes.

## INTRODUCTION

Classical CD14^+^CD16^−^ monocytes constitute about 85% of the human circulating monocyte pool. Compared to other subsets, these monocytes are adeptly prepared for host defense programs, excel in innate immune detection of pathogens phagocytosis, and display an increased capability to release cytokines (Wong *et al*, 2011). Monocyte-driven innate immune responses can yield extreme outcomes. While heightened monocyte activity triggers hyper-inflammation and cytokine storms (Bonnet *et al*, 2021; Fajgenbaum & June, 2020; Ferreira *et al*, 2021; Jafarzadeh *et al*, 2020; Junqueira *et al*, 2022; Schulte-Schrepping *et al*, 2020; Vanderbeke *et al*, 2021), dysfunctional monocytes contribute to a potentially life-threatening state of hypo-responsiveness, known as ’immunoparalysis,’ predisposing patients to opportunistic infections. Monocyte immunoparalysis is often observed after sepsis (Arens et al, 2016; Frazier & Hall, 2008; Roquilly et al, 2020), major visceral surgery (Frazier & Hall, 2008), and has recently been associated with the severity of SARS-CoV-2 infections (Agrati *et al*, 2020; Arunachalam *et al*, 2020). Monocytes are also vital cells mediating “trained immunity,” a series of long-lasting epigenetic and metabolic adaptations that enhance innate immune responsiveness upon subsequent encounters with pathogen molecules (Bekkering *et al*, 2014; Netea *et al*, 2016). Hence, understanding the mechanisms regulating monocyte functions is exceptionally relevant in diverse clinical settings, as inappropriate monocyte activities can have long-lasting immunological consequences (Agrati *et al*., 2020; Arunachalam *et al*., 2020; Bonnet *et al*., 2021; Ferreira *et al*., 2021; Jafarzadeh *et al*., 2020; Junqueira *et al*., 2022; Schulte-Schrepping *et al*., 2020; Vanderbeke *et al*., 2021).

Here, we reveal a crucial role for platelets in the effector inflammatory functions of human monocytes. In the bloodstream, monocytes continuously interact with platelets, forming monocyte-platelet aggregates (MPAs) under physiological conditions (Rinder *et al*, 1991). MPAs increased in numerous inflammatory and thrombotic disorders and are usually associated with poor outcomes (Allen *et al*, 2019; Liang *et al*, 2015; Manne *et al*, 2020; Stephen *et al*, 2013). Interaction with platelets amplifies several effector functions of monocytes in numerous diseases (D’Mello *et al*, 2017; Fu *et al*, 2021; Rong *et al*, 2014; Singhal *et al*, 2017). Nevertheless, the physiological functions of platelets or MPAs to monocyte innate functions remain undefined. Moreover, studies in the human system are lacking and were primarily done in conditions where platelets are added to monocytes or describing pathological conditions where MPAs are enhanced.

In this study, we revealed a previously unrecognized critical dependency on platelets for the cytokine production of human monocytes. Using different human monocyte isolation kits and untouched monocytes from patients with primary immune thrombocytopenia (ITP), an autoimmune disease characterized by low blood platelet counts, we demonstrate that platelet numbers directly impact the cytokine output of CD14^+^ monocytes towards Toll-like receptor (TLR) and Nod-like receptor (NLR) stimulation. Removal of platelets from healthy human monocytes caused monocyte immunoparalysis, characterized by transcriptional silencing of pro-inflammatory genes and impaired capacity to secrete pro-inflammatory cytokines. Notably, monocyte immunoparalysis is dynamic and can be reversed by replenishing monocytes with fresh platelets. Moreover, monocytes from ITP patients were inherently impaired in their capacity to produce cytokines in response to TLR and NLR stimulation. Remarkably, the supplementation of ITP monocytes with healthy platelets reactivated monocytes, reinvigorating their cytokine responses.

Mechanistically, we show that platelets are abundant cellular sources of transcription factors (TFs) and signaling molecules, including numerous NFκB and MAPK p38 subunits, which are potent cytokine synthesis regulators. Furthermore, we used Stable Isotope Labeling with Amino acids in Cell culture (SILAC) combined with high-resolution mass spectrometry (MS)-based proteomics to demonstrate the transfer of platelet-derived transcription factors (TFs), namely p65 (RelA), p52/p100 (NFκB2) and p38 MAPK into human monocytes. Finally, we reveal the capacity of fresh platelets to bypass pharmacological inhibition or genetic ablation of p38, RelA, and NFκB2 in human monocytes and restore their impaired cytokine secretion in a platelet:monocyte ratio-dependent manner. Our findings elucidate the pivotal role of platelet-derived TFs in regulating monocyte pro-inflammatory activity, suggesting the potential therapeutic application of platelet supplementation against monocyte immunoparalysis. Our research challenges the earlier notion of monocytes being immune and self-sufficient, unveiling their reliance on platelets for optimal pro-inflammatory cytokine production.

## RESULTS

### Platelets are required for cytokine secretion by human monocytes

Hyper-inflammation, resulting from inflammasome activation in monocytes, leads to excessive production of pro-inflammatory cytokines (Junqueira *et al*., 2022; Rodrigues *et al*, 2021). To thoroughly analyze the inflammasome-mediated cytokine response in human monocytes, we isolated classical monocytes (CD14^+^CD16^-^) via immune-magnetic isolation kits (**Fig S1**, and **Materials and Methods**). Consistent with previous studies (Bhattacharjee *et al*, 2017; Han *et al*, 2020; Rolfes *et al*, 2020), monocytes purified through standard magnetic isolation methods, termed “standard monocytes” (StdMo), comprised a mixture of platelet-free monocytes, monocyte-platelet aggregates (MPAs), and free platelets (Fig S**1A-D**). Employing a kit-provided platelet-depleting antibody to remove platelets from StdMo allowed us to obtain platelet-depleted monocytes (PdMo) while maintaining stable monocyte counts (Fig S**1A-C**). PdMo remained viable and were phenotypically similar to StdMo (Fig S**1A**). However, upon exposure to lipopolysaccharide (LPS) or LPS + nigericin (LPS + Nig), the secretion of IL-1β, TNFα, and IL-6 in PdMo drastically declined compared to StdMo, highlighting an essential role of platelets in monocyte cytokine production (Fig **1A-B** and S**1D-E**). Notably, reintroducing autologous platelets to PdMo (PdMo + Plts) revitalized PdMo’s impaired cytokine responses. Importantly, platelets alone (Plts) did not secrete any of the measured cytokines, confirming monocytes as their primary source. Consistent with pyroptosis downstream of inflammasome activation, we also observed reduced viability in nigericin-treated StdMo (Fig **1C**). However, platelet depletion (PdMo) led to decreased caspase-1 activity (Fig **1D**) and intracellular maturation of IL-1β (Fig **1B**) while increasing cell viability (Fig **1C**). Platelet supplementation (PdMo + Plts) restored caspase-1 activity, IL-1β maturation, and release and re-sensitized platelet-depleted monocytes to nigericin-induced pyroptosis (**Fig 1B-D**). These findings reveal the dependency on platelets for the monocyte cytokine production. To validate our findings using alternative methods, we juxtaposed two techniques for isolating CD14^+^ monocytes: positive and negative selection. We found different degrees of platelet contamination between positive vs. negative selection of CD14^+^ monocytes (Fig S**2**). Positively selected StdMo contained fewer platelets (Fig S**2B-C**) but responded poorly to TLR and inflammasome stimulation than negatively isolated StdMo, which had significantly more platelets (Fig S**2D**). The impairment of CD14-positively-selected monocytes was not due to pre-engagement of this receptor by the magnetic beads (Bhattacharjee *et al*., 2017), as these cells also displayed reduced cytokine response to Pam3CysK4, a TLR1/2 agonist. Notably, supplementing positively selected monocytes with platelets enhanced their cytokine responses (Fig S**2D**). These findings conclusively demonstrate that platelets directly impact the cytokine output of monocytes and exclude that the effects we observed were due to a specific isolation method. These findings support the requirement for platelets for inflammasome functions of human monocytes.

**Fig 1.**
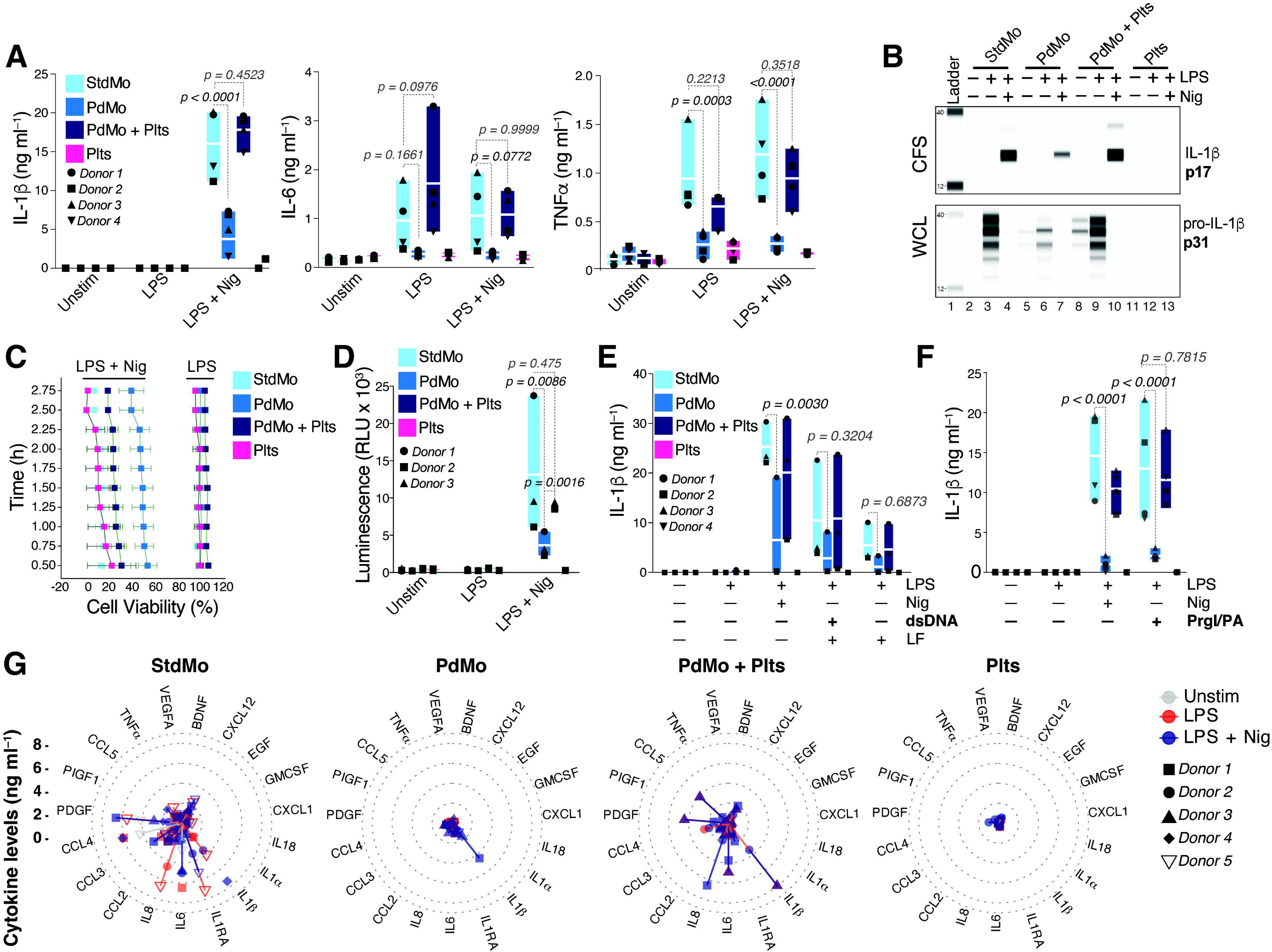
Platelets regulate the cytokine secretion of human monocytes. **(A)** Concentrations of IL-1β, TNFα, and IL-6 released into the cell-free supernatants (CFS) by untouched (StdMo, light blue), platelet-depleted (PdMo, blue), or PdMo that were supplemented with autologous platelets (PdMo + Plts, dark blue) at a 100:1 platelet:monocyte ratio. Cytokine levels secreted by platelets alone (Plts, purple) were measured as control. Cells were stimulated with LPS (2 ng ml^-1^ for 3h) followed by activation with nigericin (10 µM for 1.5 h). **(B)** Intracellular maturation and secretion of IL-1β assessed by WES capillary electrophoresis coupled with Ab-based detection in whole cell lysates (WCL) or CFS of primary human monocytes stimulated as in **A**. **(C)** Cell viability assessed every 15 min in StdMo, PdMo, PdMo + Plts, and Plts treated as in **A**. **(D)** Fluorometric assessment of caspase-1 activity in the CFS of primary human monocytes stimulated as in **A**. (**E** - **F**) IL-1β concentrations in CFS of LPS-primed StdMo, PdMo, or PdMo + Plts that were transfected with Lipofectamin (LF) containing (**E**) double-strand DNA (dsDNA, 0.5 µg ml−1) for the activation of the AIM2 inflammasome. In (**F**), cells were treated with PrgI (100 ng ml^−1^) and protective antigen (PA, 1 μg ml^−1^) for 90 min, for the activation of the NLRC4 inflammasome. Floating bars display the max/min values with indication for the mean (white bands). Each symbol represents one independent experiment or blood donor. (**G**) Radar plots displaying the 19 significantly (p < 0.05, 2-Way Anova, Tukey’s multiple comparison test) altered cytokines detected by Cytokine Luminex in the CFS of human monocytes and platelets stimulated as in **A**. Protein concentrations are represented by the spread from inner (0 ng ml^-1^) to outer circles (≥ 10 ng ml^-1^). Colors represent stimuli (Grey: Unstim, Red: LPS, dark blue: LPS + Nig). Each symbol represents one independent experiment or blood donor (n = 5 for monocytes, and n = 2 for platelets alone). For better visualisation, cytokine concentrations (y axis) are displayed with a cut off at 8 ng ml^-1^. Levels of IL-1RA and CCL4 reached 20 ng ml^-1^. **See also Fig S1 and EV2**.

In human monocyte-derived macrophages (hMDMs), platelets specifically license NLRP3 mRNA and protein expression, exclusively boosting the activation of the NLRP3 inflammasome (Rolfes *et al*., 2020). However, in human monocytes, we found that platelet removal additionally modulated the activity of the AIM2 and NLRC4 inflammasomes, indicating a broader influence of platelets on monocyte inflammasome function (Fig **1E-F**) and suggesting that platelets may influence additional innate immune sensors in monocytes. Supporting this conclusion, platelet removal impacted the monocyte production of inflammasome-independent cytokines IL-6 and TNFα upon exposure to LPS (Fig **1A** and S**1E**), Pam3CysK4, a TLR2 ligand, and R848, which activates TLR7 and TLR8 (Fig S**1F**). Platelet depletion further affected the production of numerous cytokines, chemokines, and growth factors upon TLR4 and NLRP3 inflammasome activation (Fig **1G**) or TLR1/2 and TLR7/8 activation (Fig S**1G**). While StdMo released copious amounts of IL-1β, IL-6, CCL2, CCL4, IL-8, PDGF, and IL-1RA (> 10 ng ml^-1^), the overall cytokine response of PdMo was blunted. However, the PdMo’s impaired cytokine response was reverted by reintroducing autologous platelets (50:1 platelet:monocytes ratio) to PdMo (Fig **1G** and S**1G**). Hence, the disruption of cytokine responses of PdMo was not limited to a specific TLR or the NLRP3 inflammasome but portrayed a widespread impairment of monocyte inflammation towards pattern recognition receptors (PRRs). Together, our findings underscore the paramount role of platelets in the PRR-induced cytokine response of human monocytes.

### Platelet Supplementation reverts monocyte immunoparalysis in Immune Thrombocytopenia

To assess the impact of platelets on the effector functions of monocytes in a clinical setting where blood platelet counts are inherently low, we isolated monocytes from patients with ITP. ITP is an autoimmune disorder characterized by the destruction of platelets and impaired platelet production in the absence of infections or other causes of thrombocytopenia (Cooper & Ghanima, 2019). Patients in our ITP cohort had low platelet counts despite showing average blood leukocyte counts and plasma C-reactive protein concentrations (Fig **2A-D**). Importantly, ITP patients were asymptomatic, free of infections, and were not treated with glucocorticoids (Fig **2A**). We found that untouched monocytes taken from ITP patients (referred to as “ITPMoℍ) were inherently impaired in their capacity to produce cytokines in response to LPS and LPS + Nig stimulation compared to monocytes from healthy volunteers (StdMo) (Fig **2E-G**). Remarkably, when healthy fresh platelets were supplemented to ITPMo, their cytokine production was revitalized to levels equivalent to stimulated healthy monocytes (StdMo), demonstrating that platelets can revert monocyte immunoparalysis. These findings confirm the relevance of platelets as checkpoints for monocyte-driven immune responses in a clinical setting and highlight the potential of platelet supplementation in counteracting monocyte immunoparalysis in ITP.

**Fig 2.**
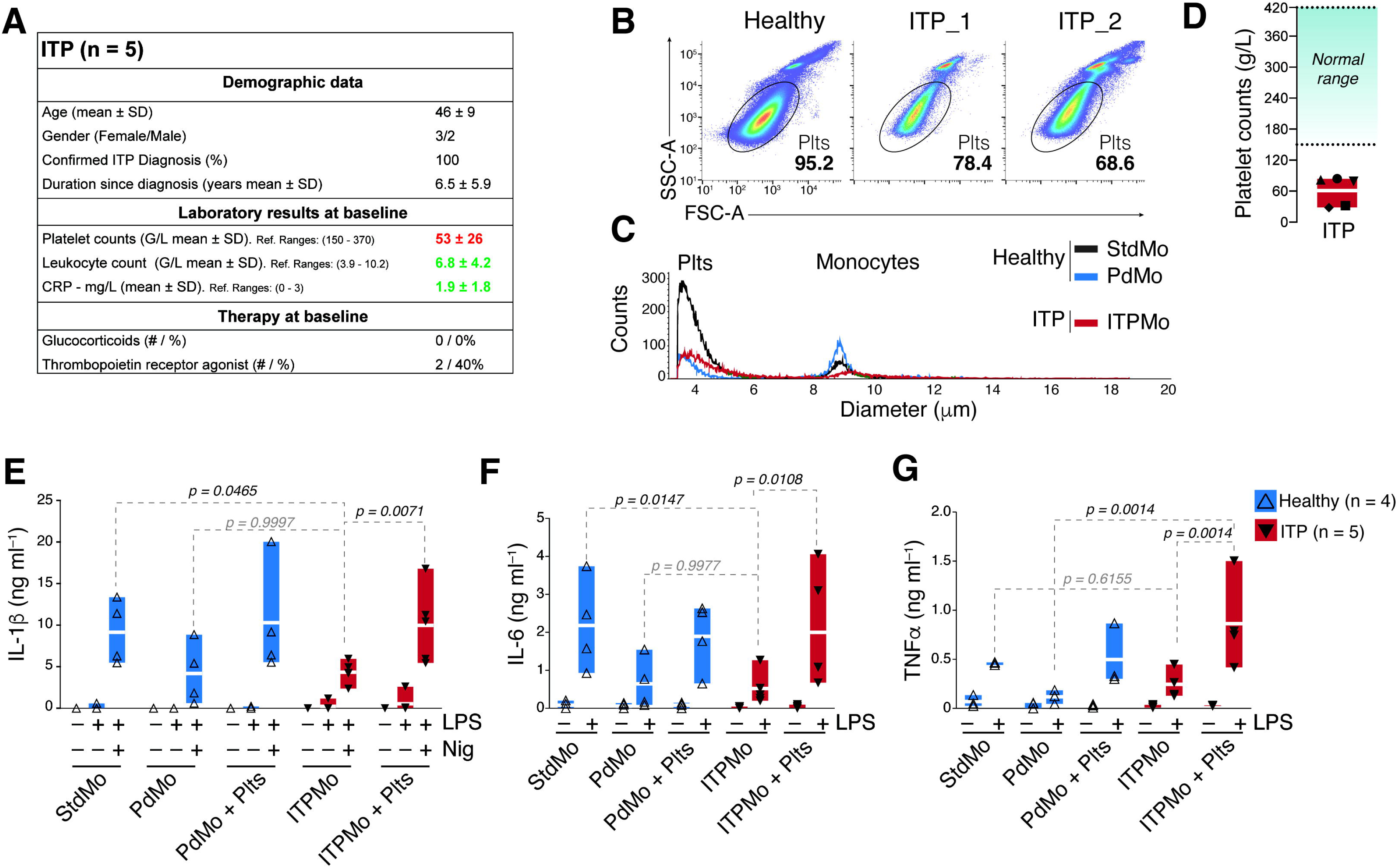
Platelet supplementation reverts monocyte immunoparalysis in Immune Thrombocytopenia (ITP). **(A)** Demographic and clinical characteristics of ITP patients (n = 5). **(B)** Representative flow cytometry percentages and (**C**) CASY automated cell quantification of monocytes and platelets in preparations of untouched or platelet-depleted monocytes from healthy donors (± platelet-depletion) or untouched monocytes from ITP patients (ITPMo). **(C)** Clinical laboratory quantification of platelets in the peripheral blood of ITP patients. Blue shaded area display the healthy reference ranges. (**E**-**G**) Concentrations of IL-1β, IL-6 and TNFα released into the CFS by untouched (StdMo), and PdMo isolated from healthy volunteers (n = 4), or untouched monocytes from ITP patients (ITPMo, n = 5). Healthy platelet-depleted monocytes (PdMo + Plts), or ITPMo were supplemented with platelets (ITPMo + Plts). Cells were stimulated with LPS (2 ng ml^-1^) for IL-6 and TNFα measurements, or LPS + Nig (10 µM) for IL-1β measurements. Data is displayed as floating bars with the max/min values and mean (white bands). P values were calculated with 2-Way Anova, Tukey’s multiple comparison test, and are displayed in the Fig. Each symbol represents one independent experiment or blood donor.

### Platelet depletion induces transcriptional silencing of inflammatory genes in human monocytes

To gain insights into the transcriptional changes in human monocytes upon platelet depletion/supplementation, we analyzed the expression of 770 genes comprising the myeloid innate immune response. We compared untouched StdMo, PdMo, and PdMo replenished with autologous platelets (PdMo + Plts) under resting conditions or upon *ex vivo* stimulation with LPS (**Fig 3**).

**Fig 3.**
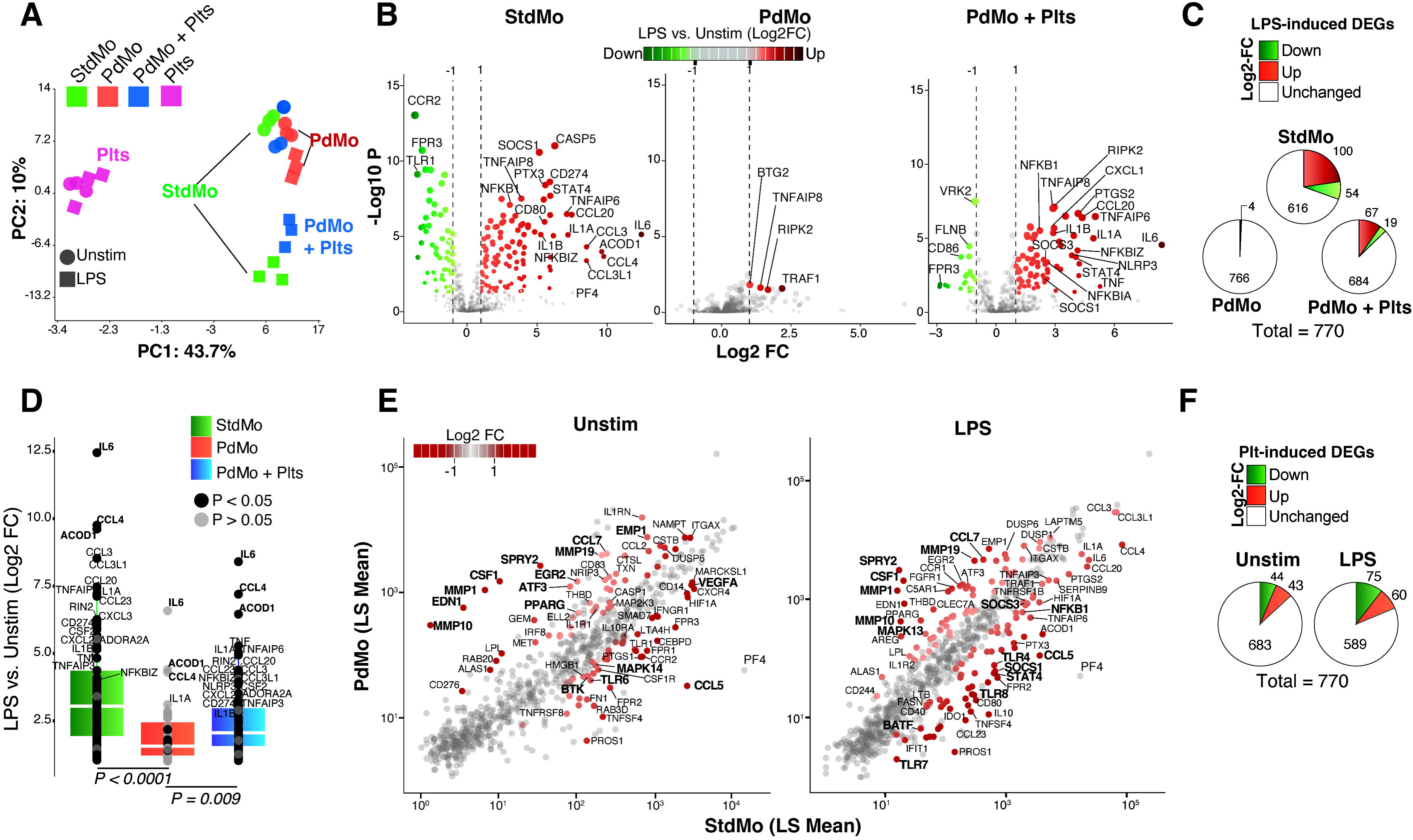
Platelet depletion causes transcriptional shutdown of myeloid innate immune genes in human monocytes. **(A)** Principal Component Analyses (PCA) of gene expression in StdMo (green), platelet-depleted monocytes (PdMo, red), platelet-depleted monocytes reconstituted with platelets (PdMo + Plts, blue), and platelets alone (Plts, pink). Each symbol represents one donor. Stimulations are represented with shapes (elypse = unstimulated, square = LPS). **(B)** Volcano plots showing log2 fold change (x-axis) and significance (−log10 *p-value; y-axis) of genes differentially expressed comparing LPS-stimulated vs. unstimulated (Unstim) StdMo, PdMo, or PdMo + Plts. **(C)** Pie charts indicate the number of up- and downregulated DEGs in each group upon LPS stimulation. **(D)** Quantitative expression (Log2 Fold Change) of the LPS-induced (>= 2 Fold Change) genes in StdMo (green), PdMo (red) or PdMo + Plts (blue). **(E)** Scatter plots displaying the LSMean (x-axis) of StdMo vs PdMo (y-axis) from DEGs comparing the effects of platelet removal in Unstim and LPS-stimulated conditions. DEGs with 2-Fold change are highlighted in red. **(F)** Pie charts representing the DEGs of the comparisons in **D**, representing DEGs induced by platelet depletion in unstimulated or LPS-stimulated monocytes. See also Fig S**3**.

Firstly, we examined the transcriptional effects of LPS or LPS + Nig stimulation in StdMo, PdMo, PdMo + Plts, and platelets alone (Plts). Principal component analysis (PCA) revealed that platelet transcripts formed distinct clusters separated from monocytes, indicating a negligible contribution of platelet transcripts to the overall mRNA pool. The transcriptional response of stimulated monocytes was clearly distinguished from that of unstimulated StdMo (**Fig 3A**), consistent with a typical LPS-induced gene expression (with 100 up-regulated and 54 down-regulated genes) compared to unstimulated conditions (**Fig 3B**, left panel). Strikingly, stimulated PdMo showed a less distinct transcriptional profile, clustering closer to unstimulated StdMo (**Fig 3A**), indicating a loss of transcriptional response to stimulation. Indeed, only four genes were significantly induced by LPS in PdMo, indicating a general suppression of inflammatory gene expression. Notably, the reintroduction of autologous platelets to PdMo (50:1 Platelet:Monocyte ratio) re-approximated their PCA-clustering towards StdMo (**Fig 3A**) and restored the expression of 86% of LPS-induced genes in PdMo, resembling the profile of StdMo (**Fig 3B**, right panel). These results were consistent with the multiplex cytokine analysis (**Fig 1G** and **S1F**).

Next, to specifically address the effects of platelet depletion/replenishment on monocytes, we separately compared StdMo vs. PdMo in unstimulated and stimulated conditions (**Fig 3E-F**). Platelet depletion alone modified 87 genes in steady-state monocytes, notably suppressing genes like MAPK14 (p38α) and BTK involved in pro-inflammatory signaling. Meanwhile, numerous transcription factors (TFs) related to monocyte differentiation and other processes were induced (Fig S**3A-C**). We identified several monocyte cytokine function regulators among the most highly differentially expressed genes (DEGs). For example, genes involved in sensing chemokines or external stimuli (e.g., *CCR2*, *FCGR3A*, and *CD14*) were down-regulated in PdMo. In contrast, several TFs (e.g., *EGR2*, *PPARG*) and ERK-MAPK repressor genes (e.g., *ATF3* and *SPRY2*) were up-regulated in PdMo (**Fig 3E-F** and S**3**). These findings demonstrate that transcriptional reprogramming underlies the functional effects of platelet depletion in primary human monocytes. Furthermore, in line with the cytokine secretion (**Fig 1G** and S**1F**), the transcriptional shutdown of pro-inflammatory gene expression in PdMos can be reversed by their replenishment with fresh platelets.

### Monocytes from thrombopenic mice display altered host immune defense transcriptional programs

Our results in ITP patients demonstrate that monocytes from thrombocytopenic individuals display naturally impaired cytokine responses to TLR and NLR activation. To recapitulate these findings *in vivo* and investigate the underlying mechanisms, we induced thrombocytopenia in mice by i.v. injection of 2 mg/kg of body weight of a rat anti-glycoprotein Ibα (GPIbα) mAb or, as control, with equal amounts of a rat IgG isotype (Sreeramkumar *et al*, 2014; Xiang *et al*, 2013) (Fig **4A**). We then challenged platelet-depleted mice with i.v. injection of LPS (2 mg/kg). Antibody-dependent platelet clearance efficiently lowered blood platelet counts in treated mice (Fig **4B**). Contrary to what our experiments with human monocytes indicate, but in line with earlier findings in mice injected with LPS or in models of bacterial infection (Carestia *et al*, 2019; Claushuis *et al*, 2016; de Stoppelaar *et al*, 2014; Xiang *et al*., 2013), platelet depletion did not significantly influence the plasma levels of IL-1β, TNFα, and IL-6 in challenged mice (Fig S**4A**).

**Fig 4.**
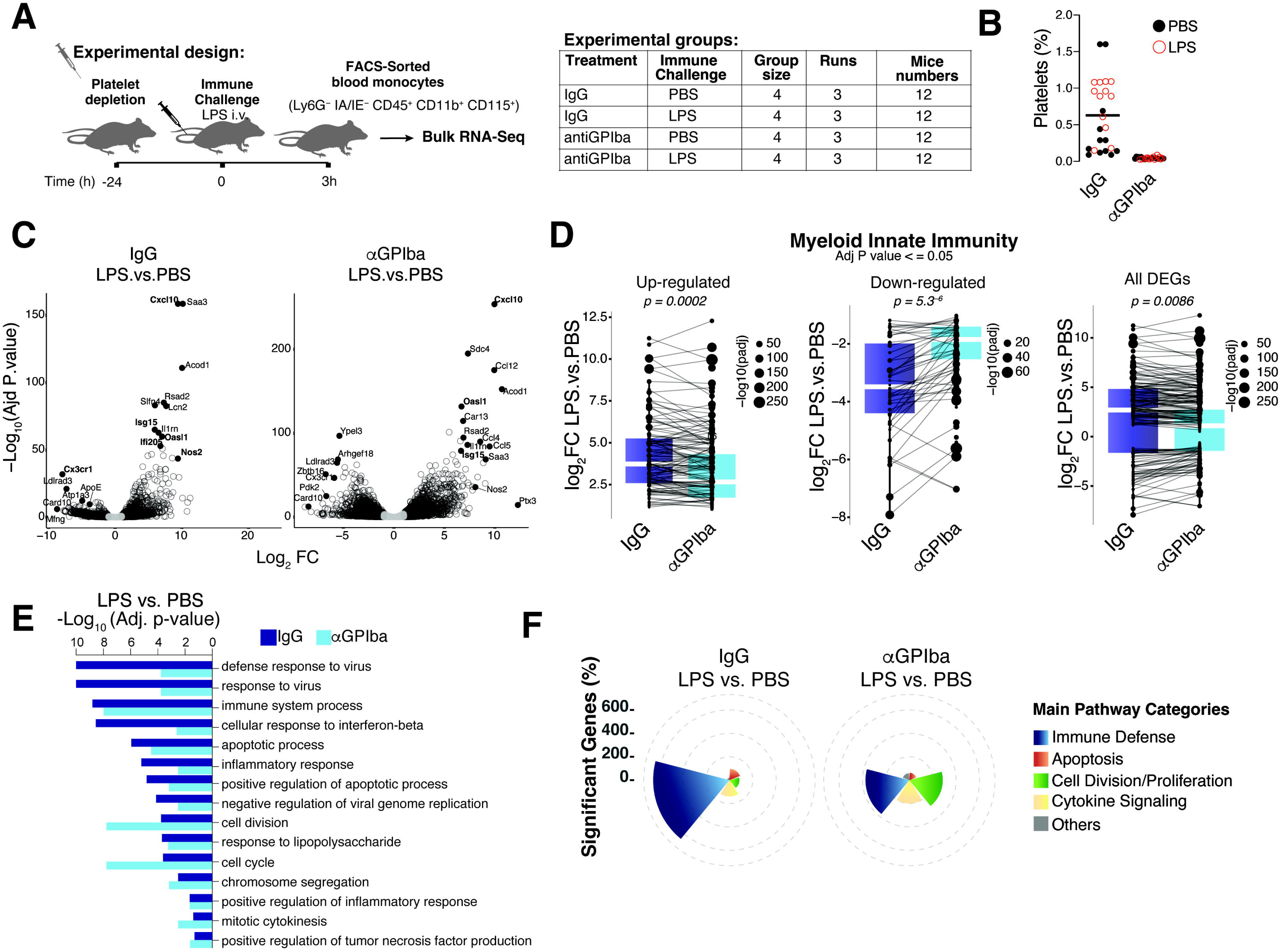
Dysregulation of host defense transcriptional programs in monocytes from thrombopenic mice. **(A)** Schematic representation of the experimental setting for platelet depletion in vivo depletion followed by intravenous challenge with LPS, FACS-sorting and bulk-RNASeq of blood monocytes. Monocytes were isolated from PBMCs from pooled blood of 4 mice per group (IgG- or anti-GPIbα-treated) each challenged either with LPS or PBS in 3 different experiments. **(B)** Flow cytometric quantification of platelets in IgG-treated and anti-GPIba treated mice (n = 24 per condition). **(C)** Volcano plots (log2 fold change vs. −log_10_ adjusted P-value) of the significant DEGs (adjusted P-value < 0.05, Fold Change ≤ −2 or ≥ 2) comparing PBS vs. LPS-challenge in IgG (left) or anti-GPIba-treated mice (right). **(D)** Fold change of the DEGs that compose the myeloid innate immunity that were modulated by LPS in monocytes from IgG vs. antiGPIbα-treated mice. P values from t.test are shown. **(E)** Pathway enrichment analysis of the LPS-induced DEGs comparing monocytes from IgG (dark blue) or anti-GPIba-treated mice (light blue). **(F)** Main represented categories of pathways (GO analysis) from the DEGs induced by LPS on monocytes from IgG-treated vs. platelet-depleted mice (See also Fig S**4**).

To investigate the effects of platelet depletion specifically on blood monocytes, we performed RNASeq analysis on FACS-sorted (Ly6G^−^ IA/IE^−^ CD45^+^ CD11b^+^ CD115^+^) murine PBMCs. Consistent with the removal of platelets by anti-GPIbα mAbs, the expression of several platelet transcripts (e.g., *Pf4*; *Gp9*; *Itga2b*, *Ppbp*, *Tubb1*, *Treml1,* and *Clu*) was down-regulated in monocytes from platelet-depleted compared to IgG-treated mice in the absence of immune stimulation (PBS group) (Fig S**4B**). Nevertheless, platelet depletion increased the expression of genes involved in the complement cascade (e.g., *C1qa*, *C1qb,* and C1qc) and cell activation processes (e.g. *Htra3* and *Mertk*) (Fig S**4B**), correlating with enrichment in pathways involved in blood coagulation, hemostasis, platelet activation, complement, and wound healing processes (Fig S**4C**), and suggesting that anti-GPIbα-mediated platelet depletion resulted in platelet activation.

We next compared the effects of platelets on the monocyte transcriptional response to LPS challenge *in vivo*. Expression profiling analysis (Fig **4C**) and gene ontology (GO) (Fig **4E-F**) confirmed that monocytes isolated from IgG-treated mice displayed a typical LPS-induced transcription profile (Alasoo *et al*, 2015) with the enrichment of genes involved in immune defense among the highest DEGs (e.g. *Cxcl10*, *Ccl12, Il1a, Ly6c,* interferon-stimulated genes (ISGs), and *Il12a*) and the concomitant and expected down-regulation of *Card10 and Cx3cr1* (Pachot *et al*, 2008) (Fig **4C**). Overall, LPS stimulation altered the expression of a higher number of transcripts in monocytes from platelet-depleted (anti-GPIbα-treated) compared to IgG-treated mice (4088 vs. 1282 transcripts, Fig S**4D**). However, the expression of several LPS-induced genes was comparable between anti-GPIbα and IgG-treated mice (Fig S**3E**). More than 80% of the LPS-induced gene signature was shared between monocytes from IgG-treated and platelet-depleted mice. While 3008 DEGs were exclusively expressed in monocytes from platelet-depleted mice, 202 DEGs were exclusively regulated in monocytes from IgG-treated mice (Fig S**4D**). Next, we focused the analysis to encompass the 770 myeloid innate immunity genes used in the human monocyte NanoString (**Fig 3**). Akin to the transcription profile of human PdMos, monocytes from thrombopenic mice had a significantly reduced expression of genes related to the myeloid innate immune response in response to i.v. LPS challenge (Fig **4D**). Consistent with the importance of the myeloid compartment in innate immunity, the LPS-induced expression profiles identified higher enrichment scores for pathways involved in host immune defense programs in IgG-treated mice (e.g., immune response to virus, immune system processes, cellular responses to IFNβ, and inflammation (Fig **4E-F**). These pathways were also represented but with lower enrichment scores in the gene profile of platelet-depleted mice, which was skewed towards apoptosis and cell cycle regulation (Fig **4E-F**). From these findings, we conclude that, although monocytes in thrombopenic mice maintain their proinflammatory cytokine capacity, platelet depletion induces transcriptional reprogramming towards cell cycle and division at the expense of immune defense programs. These findings align with previous observations of compromised immunity (Carestia *et al*., 2019; Claushuis *et al*., 2016; van den Boogaard *et al*, 2015; Xiang *et al*., 2013), and higher susceptibility to sepsis in thrombopenic mice, despite normal inflammatory responses.

### Platelets regulate cytokine secretion in human monocytes in trans and independently of classical platelet-monocyte crosstalk mechanisms

Evidence highlights that innate immune cells can engulf platelets (Lang *et al*, 2002; Maugeri *et al*, 2009; Rolfes *et al*., 2020; Senzel & Chang, 2013). Our observations did not reveal distinguishable internalization of platelets by human monocytes (Fig **5A**). However, blocking actin polymerization in platelet-depleted monocytes (PdMo) via Cytochalasin D (CytoD) (Fig **5B**) or Latrunculin B (Lat-B) (Fig S**5A**) ablated the ability of platelets to restore their faulty cytokine secretion. The potential role of endocytosis in this mechanism was also assessed, but the treatment of PdMo with Dynasore (DS), an inhibitor of dynamin 1/2 endocytosis, did not prevent the platelets’ ability to reconstitute the faulty cytokine response of PdMo (Fig **5B**). These data underpins the significance of monocyte’s phagocytic machinery for the influence exerted by platelets, albeit without directly visible platelet phagocytosis, supporting a scenario where platelet-derived vesicles (PMPs), although too small for imaging, are internalized by monocytes (Vajen Mause & Koenen, 2015). Supporting this conclusion, platelet released molecules (Plts Sups) efficiently restored cytokine production in LPS-stimulated human PdMo (Fig **5C**).

**Fig 5:**
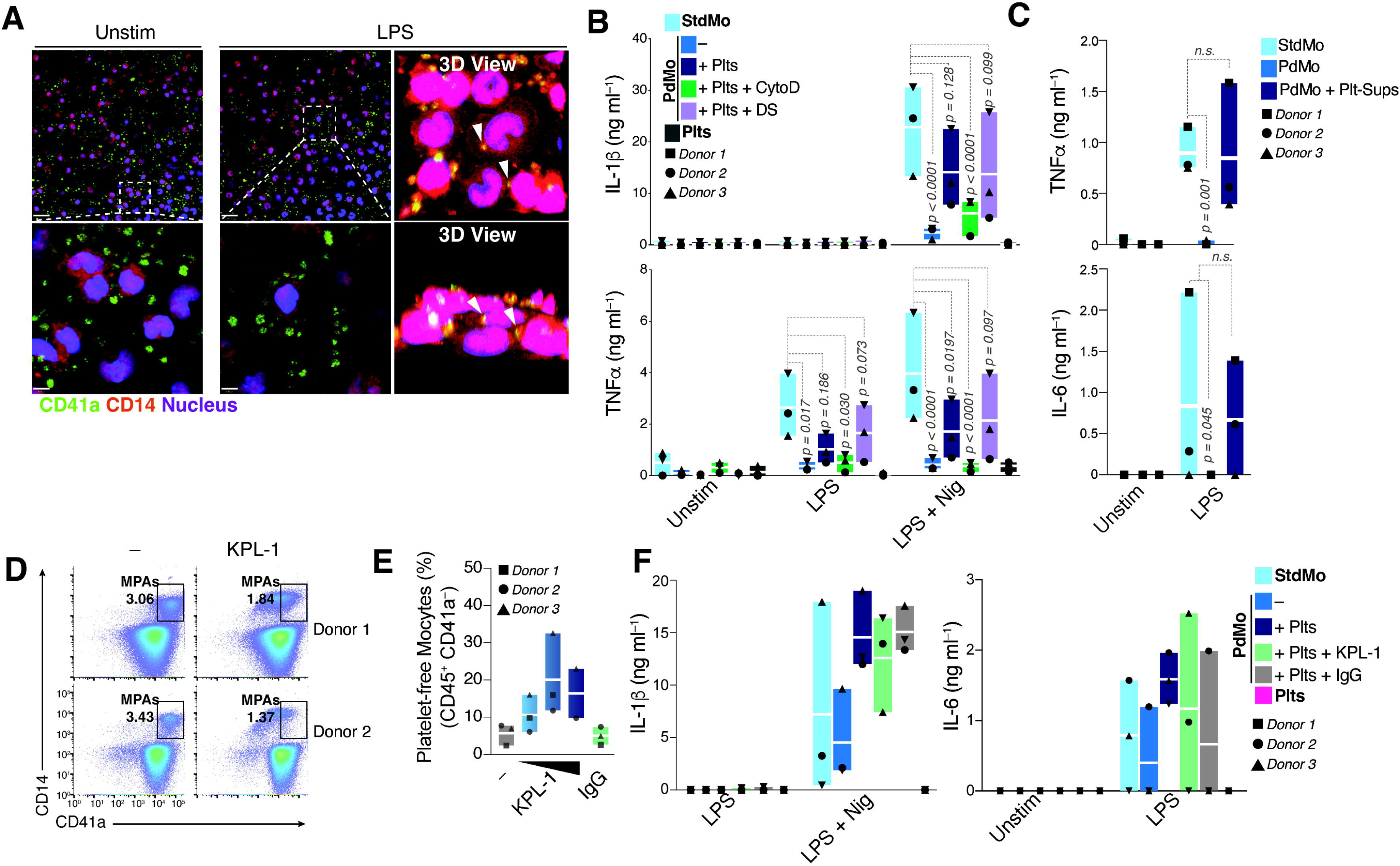
Platelets modulate monocyte cytokine responses in trans. **(A)** Confocal imaging of StdMo and platelets. Cells were stained with CD41a AF488 (platelets: green) and CD14 AF647 (monocytes: red). Nuclei were stained with Hoechst 34580 (blue). Scale: 24 μm (top panel); 4.8 μm (bottom panel). White arrows indicate points of contacts between platelets and monocytes. Images are from one representative of four independent experiments. **(B)** IL-1β and TNFα levels in CFS of StdMo, PdMo, and PdMo that were supplemented with autologous platelets (PdMo + Plts, 100:1 platelets/ monocytes). Monocytes were pre-treated with Cytochalasin D (CytoD, 50 μM) or Dynasore (DS, 30 μM) before being supplied with Plts. Cells were stimulated with LPS (2 ng ml-1), followed by activation with nigericin (10 μM). **(C)** TNFα and IL-6 levels in CFS of LPS stimulated StdMo, PdMo, PdMo, or PdMo reconstituted with platelet releasates (Plt-Sups). **(D)** Representative flow cytometry assessment and gating strategy of StdMo that were incubated with growing concentrations of KPL-1 (5, 10 or 25 μg ml^-1^), or IgG (25 μg ml^-1^). Gates indicate the frequencies of free platelets (CD45- CD41a+), MPAs (CD45+CD41a+) and platelet-free monocytes (CD45+ CD41a-). Data is representative of two independent experiments with several donors. **(E)** Cumulative flow cytometry frequency of free monocytes (CD45^+^ CD41a^-^) in StdMo treated with growing concentrations of KPL-1 (5, 10 or 25 μg ml-1), a specific mAb against P-selectin, or an unrelated isotype IgG control (25 μg ml^-1^). See **Fig S5B** for gating strategy. **(F)** IL-1β, TNFα and IL-6 levels released by stimulated StdMo that were treated with increasing concentrations of KPL-1 (5, 10 or 25 μg ml^-1^), or IgG (25 μg ml^-1^).

Lastly, we examined whether ligand-receptor interactions mediate platelet-monocyte crosstalk. A well-documented mechanism of monocyte-platelet or monocyte-PMP binding hinges on the interaction between platelet P-selectin (P-sel or CD62P) and P-selectin glycoprotein ligand-1 (PSGL-1 or CD162) present on monocytes (Frenette *et al*, 2000; Han *et al*., 2020). To delineate the importance of the Psel-PSGL-1 axis, we employed KPL-1, an antibody targeting P-sel. Although KPL-1 markedly reduced monocyte-platelet binding (**Fig 5D-E** and S**5B**), it neither inhibited pro-inflammatory cytokine production in stimulated StdMo nor impeded platelets in restoring cytokine dysfunction in PdMos (**Fig 5F**). Although both soluble and immobilized P-sel can influence innate immune functions in stimulated monocytes (Weyrich *et al*, 1995), and platelets can release P-sel in vesicles (Forlow McEver & Nollert, 2000), introducing soluble recombinant human P-sel to PdMo didn’t rectify their impaired cytokine secretion, compared to the re-addition of autologous platelets (**Fig S5C**). These insights suggest the P-sel-PSGL-1 interaction is not central to revitalizing the faulty cytokine response of PdMo. Despite KPL-1 causing a physical separation between platelets and StdMo, it did not hinder cytokine secretion, supporting the involvement of vesicles that may still be present.

As the platelet-monocyte crosstalk is additionally facilitated by platelet glycoprotein-Ib (GPIIb) and monocyte CD11b (Carestia *et al*., 2019; Malehmir *et al*, 2019), we blocked this ligand-receptor axis with anti-CD11b, or the GPIIb/IIIa inhibitor trihydrochloride (GR-144053) (Fig S**5D**). Notably, neither targeting the GPIIb-CD11b complex nor inhibiting GPIIb/IIIa prevented the platelet’s ability to rescue the faulty cytokine production in platelet-depleted monocytes (Fig S**5D**). Next, we blocked integrins, additional players in platelet adhesion, and interactions with leukocytes (Bennett Berger & Billings, 2009; Nieswandt *et al*, 2000). We used RGD, an antagonist of platelet glycoprotein IIb/IIIa receptors and other integrins (Haskel & Abendschein, 1989). We pre-incubated platelets with RGD before their addition to PdMo. Irrespective of RGD, platelets similarly rescued cytokine secretion of PdMo (Fig S**5E**), indicating that this class of integrins is not involved in the platelet regulation of monocyte cytokine responses. Using similar inhibition strategies with mAbs, and simulation of ligands with recombinant human proteins, we further excluded a role for RANTES (CCL5) (Alard *et al*, 2015), CXCL12(Chatterjee *et al*, 2015), sialic acids (SAs)(Kullaya *et al*, 2018), Siglec-7(Varchetta *et al*, 2016), and the CD40-CD40L axis(Henn *et al*; Inwald *et al*, 2003) in the effects described in our study (Fig S**5F-H**). Together, our findings indicate that the classical and well-described mechanisms of platelet-monocyte crosstalk (i.e., co-stimulatory molecules, integrins, CD40-CD40L axis, CCL5, CXCL12, and SAs) are not involved in the effects described in our study.

### Platelets are cellular sources of transcription factors (TFs)

Our transcriptomic approach revealed that platelets influence the expression of genes controlled by key transcription regulators orchestrating pro-inflammatory cytokine production in monocytes, such as NFκB, BTK, and MAPKs. We, therefore, profiled the activation of serine/threonine (STK) and protein tyrosine (PTK) kinase networks on StdMo, PdMo, or PdMo + Plts as well as on platelets alone (Plts) and observed that platelet depletion markedly impacted the kinome network of LPS-stimulated monocytes (Fig **6A-D** and S**6A**). With few exceptions, the LPS-induced phosphorylation of numerous PTK and STK subtracts was significantly lower in PdMos compared to StdMo (Fig **6C-D**). Substantiating the results of mRNA expression (**Fig 3**), platelet depletion decreased the phosphorylation of BTK, p38 MAPK, MAPK-activated protein kinases (MAPKAPK), IKK kinases, erythropoietin-producing human hepatocellular receptors (Eph), Cyclin-depended kinases (CDK), protein kinase A, (PKA) and protein kinase C (PKC). In contrast, platelet-depleted monocytes displayed increased activation of Janus kinases (JAKs) and IKKs, which are known modulators of NFκB signaling (Solt & May, 2008) (Fig **6A**). In line with the altered function of these groups of kinases, pathway analysis revealed that platelet depletion predominantly changed the activity of kinases involved in signal transduction associated with NFκB activation, immune responses via CD40 signaling, MAPK, and JAK/STAT signaling pathways (Fig **6E**). Combined, the transcriptome (Fig **3**) and kinome (Fig **6**) of human monocytes subjected to platelet depletion/re-addition revealed dysregulation of key cytokine-regulatory pathways in PdMo.

**Fig 6.**
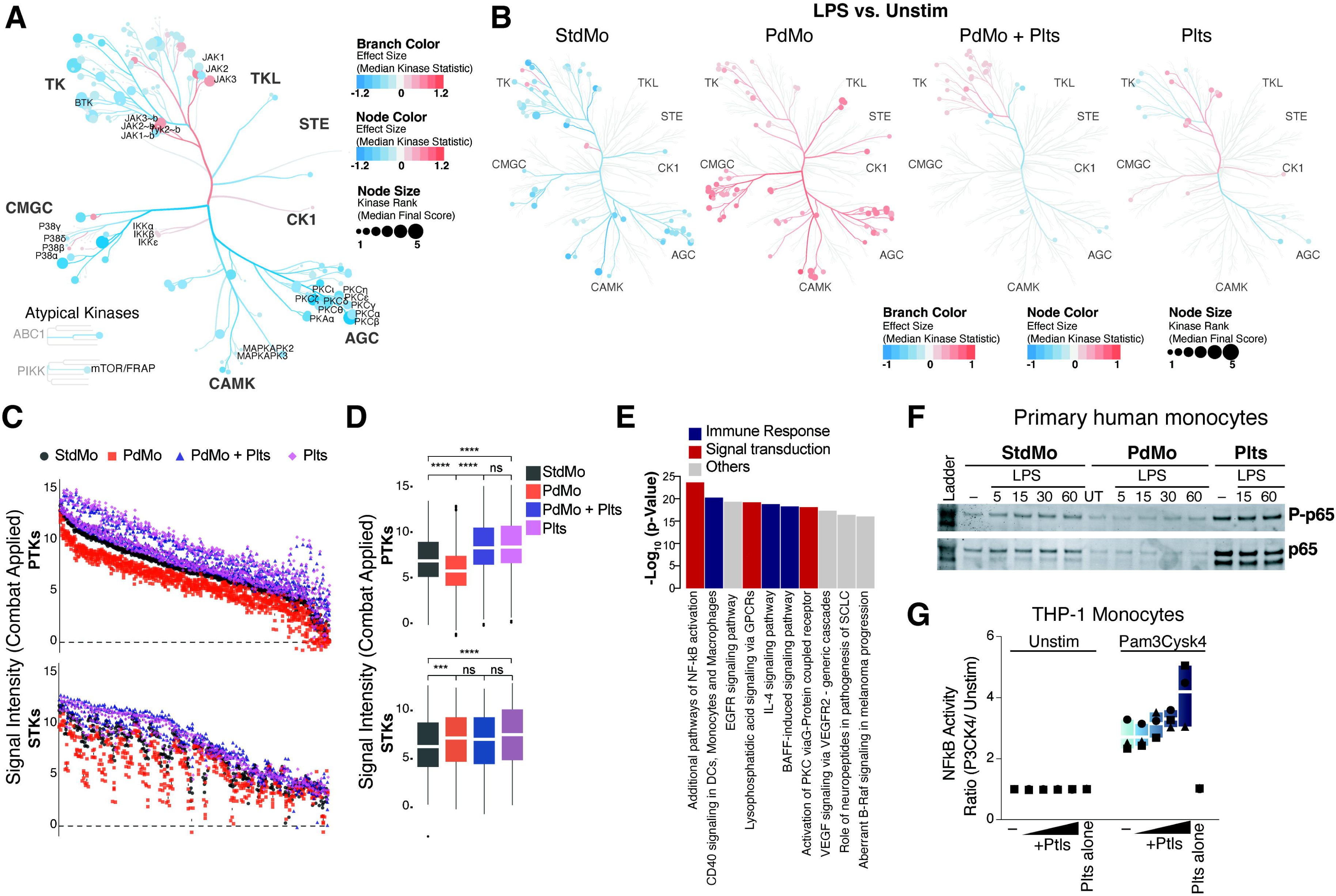
Platelets are cellular sources of active kinases and transcription factors. (**A-B**) Coral trees display the activity of Protein Tyrosine (upper branches) and Serine/Threonine kinases (lower branches) in StdMo, PdMo, PdMo + Plts and platelets from 5 donors, upon LPS stimulation for 15 min. (**A**) Impact of platelet depletion on the activity of PTK and STK in unstimulated monocytes. (**B**) Effects of LPS stimulation on the activation of PTKs and STKs in StdMo, PdMo, PdMo + Plts and platelets. Red: increased kinase activity, Blue: reduced kinase activity. (**C-D**) Scatter plots and **D** box plot showing the signal intensity for the phosphorylation status of phosphosites (x.axis) that are substracts of PTK (top) and STK (bottom) comparing StdMo, PdMo, PdMo + Plts, and platelets alone (Plts). Each symbol shows one donor (n = 5) and only include peptides which passed the QC. batch correction with a 2-step Combat correction. P values in D were calculated by Anova multiple comparison. (**E**) Pathway analysis of the most represented signaling pathways dynamically changed upon platelet depletion/re-addition to primary human monocytes. See also **Fig S6**. (**F**) Immunoblot of total RelA (p65), and Phosphorylated-p65 (P-p65) in MoStd, MoPD and Plts that were left untreated (−) or stimulated with LPS for 5, 15, 30 or 60 min. Representative of three independent experiments. (**G**) NFκB activity assay in CFS of unstimulated (Unstim), or Pam3CysK4-stimulated (100 ng ml^-1^) NFκB-SEAP and IRF-Lucia luciferase Reporter Monocytes cultured alone (−) or co-cultured with increasing concentrations of human platelets (1:10, 1:20, 1:50 and 1:100). SEAP-Activity was measured with QuantiBlue Buffer. Graphs show floating bars display max/min values with indication to the mean (white bands). Each symbol represents one donor, or independent experiment.

Confirming the dynamic changes in NFκB activation caused by platelet depletion/reconstitution in human monocytes, the LPS-induced phosphorylation of the NFκB subunits p55 (RelB) and p65 (RelA) was impaired in PdMo, as measured by HTRF (Fig S**6B**) and immunoblotting (Fig **6F**), and it was restored by the re-addition of platelets. Supporting that platelets regulate the NFκB activity on monocytes, increasing concentrations of platelets enhanced TLR2-triggered NFκB activity in THP-1 monocytes encoding an NFκB-inducible promoter (Fig **6G**). Together, these findings highlight the involvement of the NFκB and MAPK signaling pathways as candidate mechanisms by which platelets regulate the cytokine production of human monocytes.

Interestingly, throughout our kinase and immunoblotting assays, platelets consistently showed high basal kinase activity (Fig **6C-D**), and protein levels of NFκB subunits (RelA, p65) (Fig **6F** and S**6C**), and p38 MAPK (Fig S**6D**), supporting prior findings that platelets are cellular sources of transcription factors (TFs) and signaling molecules (Lannan *et al*, 2015). Indeed, by assessing the native and phosphorylated form of these proteins in the lysates or supernatants of human platelets, we demonstrated that platelets express and release NFkB and confirmed previous observations that they release p38α (Fig S**6C-D**), the major isoform expressed on human and mouse platelets (Shi *et al*, 2017). Although phosphorylated p65 (P-p65) and P-p38 were predominantly enriched in platelets, these proteins were additionally detected in platelet supernatants (Fig S**6D**) (Lannan *et al*., 2015).

### Platelets transfer TFs to monocytes

Given the abundance of these critical transcriptional regulators in non-nucleated platelets (Beaulieu & Freedman, 2009; Ezumi Takayama & Okuma, 1995; Lannan *et al*., 2015; Poli *et al*, 2021) and the corresponding dynamic activation of these pathways on platelet-depleted/replenished monocytes (**Figs 3** and **6**), we speculated that platelets transfer TFs to monocytes. Cell-to-cell propagation of TFs and other signaling molecules has been reported to occur through gap junctions between connecting cells, with biological functions demonstrated in recipient cells (Ablasser *et al*, 2013; Kasper *et al*, 2010). However, little is known about the trans-cellular propagation of signaling molecules when circulating cells adhere to one another.

To precisely delineate the repertoire of platelet proteins transferred to monocytes, we employed an unbiased proteomic approach combined with Stable Isotope Labeling by Amino acids in Cell culture (SILAC). For this, we cultured human megakaryocytes (MEG-01 cells) for several days in a medium containing stable isotope labeled amino acids C^13^L^15^] L-lysine and [C^13^L^15^] L-Arginine (hereafter referred to as heavy AAs). Under these conditions, newly translated proteins in MEG01 incorporate heavy AAs and can be distinguished from pre-existing proteins on monocytes by a mass shift (Wolf *et al*, 2020). Of note, cell-free supernatants from MEG-01 (termed MK-Sups), akin to platelets, were equally capable of restoring the impaired cytokine responses observed in PdMos (Fig **7A**). Subsequently, PdMos were incubated with MK-Sups, followed by a stringent washing protocol to eliminate non-specific associations before subjecting the monocyte lysates to MS analysis (Fig **7B-C**). We detected 33 proteins harboring heavy AAs enrichment in PdMos exposed to MK-Sups under LPS-stimulated conditions (Fig **7D**) and numerous TFs in unstimulated monocytes (Fig S**7A**). Our data revealed notable entities among these proteins, including the transcriptional activator NFκB2 (p100/p52), transcriptional regulators (e.g., ANP32B, THOC1), enzymes involved in mRNA translation and stability (ZFP36, EOF2A, and PAIP1), and proteins implicated in the MAPK/ERK signaling cascade (Fig S**7**). Furthermore, 5 out of the 33 heavily labeled proteins enriched in PdMo after LPS were involved in transmembrane functions, pointing towards a mechanism for the internalization of MK-derived proteins. We confirmed the expression and release of NFκB2 on human platelets by immunoblotting (Fig **7E**). These findings substantiate the transfer of platelet-originated transcriptional and signaling molecules to monocytes, underscoring a potentially crucial intercellular communication mechanism.

**Fig 7:**
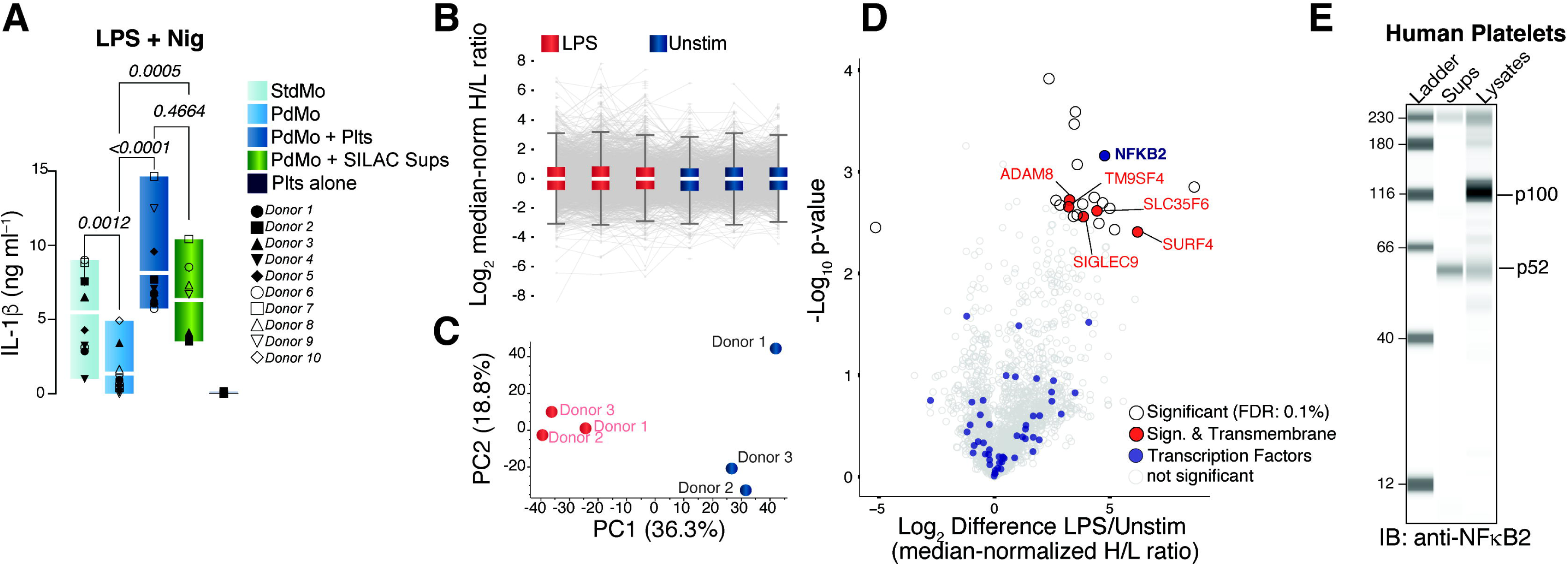
Enrichment of megakaryocyte-derived NFκB2 in human monocytes. **(A)** IL-1β released from LPS-primed StdMo, and PdMo that were reconstituted with platelets (PdMo + Plts) or culture supernatants from MEG-01 cells that were cultured in SILAC medium. Floating bars display the max/min values with indication for the mean (white bands). Each symbol represents one independent blood donor (n = 10). (**B - D**) Analysis of Mass speck proteomics combined with Stable isotope labeling with amino acids in cell culture (SILAC) in human PdMo exposed to cell-free supernatants of MEG-01 cells saturated with heavy AAs. Boxplots of quantitative distributions and (**B**) Principal component analysis (PCA) of SILAC proteomics samples from PdMo reconstituted with cell-free supernatants of MEG-01 cells. (**D**) Volcano plots of proteins with heavy-AAs detected in PdMo exposed to cell-free supernatants of MEG-01 cells through Mass speck proteomics combined with Stable isotope labeling with amino acids in cell culture (SILAC). Cells were stimulated with LPS. Unstimulated conditions are shown in **Fig S7**. (**E**) WES capillary electrophoresis and immunoblotting of NFkB2 on whole cell lysates (Lysates) or cell-free supernatants (Sups) from human platelets. Representative of 2 independent experiments.

### Platelet-derived TFs license monocyte innate immune responses

Next, to specifically address whether platelet-derived NFκB or p38 functions in monocytes, we tested whether pharmacological inhibition of NFκB and p38 MAPKs on platelets interfere with their capacity to reactivate cytokine production in PdMos (**Fig 8A**). Indeed, pre-treatment of platelets with the irreversible inhibitor of IKKβ phosphorylation BAY 11-7082 (BAY) or the p38 inhibitor SB203580 (Yamashita Hishinuma & Shimada, 2009) extinguished their ability to recover cytokine response of PdMo (**Fig 8B**). The addition of intact platelets, but not NFκB- or p38-inhibited platelets, restored IL-6 and TNFα secretion of LPS-stimulated PdMo as well as LPS + nigericin-induced IL-1β production (**Fig 8B**). To account for inhibitor carryover, we separately treated monocytes with BAY or SB203580 (**Fig 8B**, last two right bars). BAY blocked the thrombin-induced P-selectin expression on treated platelets (Fig S**8A-B**). While BAY completely abrogated monocyte cytokine responses to TLR and NLRP3 activation, direct incubation of monocytes with SB203580 partially inhibited their response. These findings support that p38 and NFκB or their downstream signaling in platelets are essential for the platelet regulation of cytokine responses of human monocytes.

**Fig 8:**
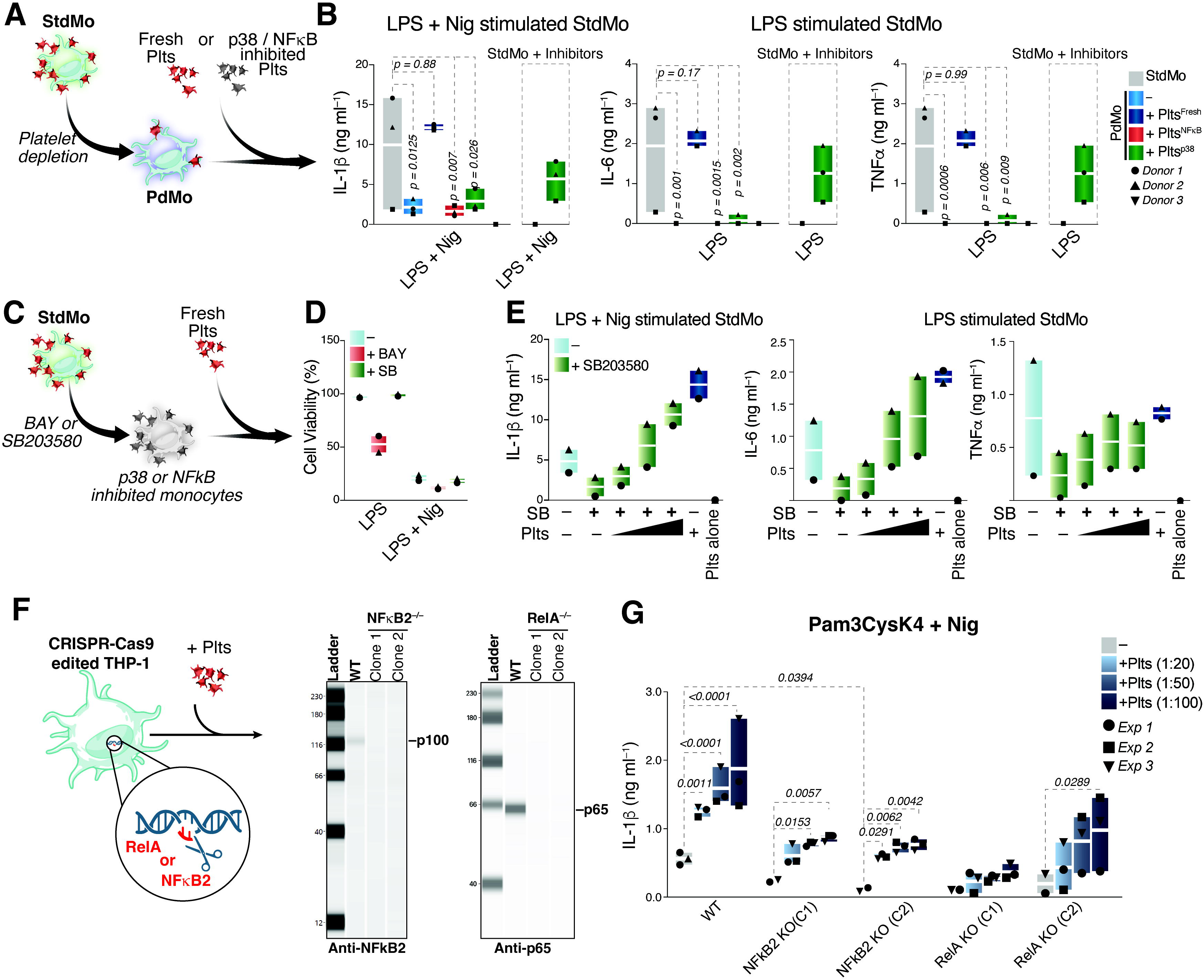
Platelets restore cytokine function in NFκB2-deficient monocytes. **(A)** Schematic representation of the supplementation of platelet-depleted (PdMo) primary human monocytes with autologous platelets that were left untreated (+ Fresh Plts), or pre-incubated for 20 min with 50 μM of BAY 11-7082 to inhibit NFκB (+ Plts^NFκB^) or 20 μM of SB203580 to inhibit p38 MAPKs (+ Plts^p38^) and washed before being added to PdMo. **(B)** IL-1β, TNFα, and IL-6 concentrations in CFS of StdMo, PdMo, or PdMo that were supplemented with platelets pretreated with BAY11-7082 (+ Plts^BAY^), SB203580 (+ Plts^SB^), or left untreated (+ Plts^Intact^). Co-cultures were stimulated with LPS (2 ng/ml) followed by nigericin stimulation (10 μM). Side bars show the effects of direct exposure of monocytes to the inhibitors (StdMo + Inhibitors). **(C)** Schematic representation of the inhibition of NFkB or MAPK p38 on primary human monocytes (StdMo) followed by their replenishment with fresh autologous platelets. **(D)** Cell viability assay in human monocytes treated with BAY 11-7082 (50 μM, 30 min) or SB203580 (20 μM, 30 min) followed by stimulation with LPS or LPS + Nig, as depicted in **C**. **(E)** IL-1β, TNFα, and IL-6 concentrations in the CFS of StdMo that were treated with SB203580 before been added with increasing ratios of freshly isolated platelets (1:5, 1:50, and 1:100). Cells were stimulated with LPS or LPS and nigericin (LPS + Nig). Floating bars display max/min values with indication to the mean (white bands). Each symbol represents one donor. See also Fig S**8**. **(F)** Schematic representation of the genetic ablation of p65 (RelA) or p52/100 (NFκB2) on human THP-1 monocytes, and knockout confirmation at the protein level by immunoblotting for each protein. Two clones are shown. Data is representative of 3 independent experiments. **(G)** IL-1β concentrations in the CFS of stimulated RelA^−/−^ or NFκB2^−/−^ THP-1 monocytes that were cultured alone or added with increasing ratios of freshly isolated platelets (1:20, 1:50, and 1:100). Cells were stimulated with Pam3CysK4 or Pam3CysK4 and nigericin (LPS + Nig). Floating bars display max/min values with indication to the mean (white bands). Each symbol represents one donor. Data is pooled from 2 independent experiments.

Next, to address whether platelet-derived TFs function in recipient monocytes, we examined whether supplementation with fresh platelets could bypass NFκB or p38 inhibition in primary human monocytes. To demonstrate this, we pre-incubated monocytes with high doses of BAY (50 µM) and SB203580 (20 µM). After washing the inhibitors away, we supplemented BAY- or SB203580-treated monocytes with growing concentrations of intact platelets freshly isolated from the same donors (Fig **8C**). In line with previous observations of toxicity associated with BAY-11-7082 in human cells (Rauert-Wunderlich *et al*, 2013; White & Burchill, 2008), BAY-treated monocytes displayed decreased viability (Fig **8D**), precluding us from addressing the importance of platelet-derived NFκB in these experiments. Consistent with dysfunctional p38 signaling (Lee *et al*, 1994), SB203580-treated monocytes were incapable of secreting cytokines in response to LPS or LPS + Nig (Fig **8E**). Strikingly, the addition of intact platelets dose-dependently bypassed p38 inhibition and restored cytokine secretion in SB203580-treated monocytes (Fig **8E**). Hence, these findings support that platelet supplementation restores dysfunctional p38 activity in p38-inhibited monocytes in a dose-dependent manner. Supporting this conclusion, SB203580-treated platelets failed to convert cytokine responses in SB203580-treated monocytes (Fig S**8C**).

Finally, to specifically address the function of platelet-derived NFκB on monocytes and to simulate a scenario in which platelets would be the only cellular source of functional NFκB, we genetically ablated RelA or NFκB2 in THP-1 monocytes using CRISPR-Cas9 (Fig **8F**). Again, and consistent with a genetic deficiency in these molecules, confirmed by immunoblotting (Fig **8F**), RelA^−/−^ or NFκB2^−/−^ monocytes were unable to secrete IL-1β in response to Pam3Cysk4 + Nig stimulation (Fig **8G**). As previously reported (Rolfes *et al*., 2020), platelets boosted IL-1β secretion in wild-type monocytes (Fig **8G**). Notably, adding platelets to RelA^−/−^ or NFκB2^−/−^ monocytes dose-dependently boosted their production of IL-1β, demonstrating the reactivation of these pathways in otherwise knockout recipient cells. These findings demonstrate that platelets supply human monocytes with functional signaling TFs, which license their optimal cytokine responses and reveal a novel layer of cell-to-cell communication in innate immunity.

## DISCUSSION

Monocytes play a pivotal role in the innate immune system, acting as both frontline defenders against pathogens, being progenitors of macrophages and dendritic cells, and releasing pro-inflammatory cytokines like IL-1β, IL-6, and TNFα crucial in initiating the inflammatory response (Guilliams Mildner & Yona, 2018). Unrestricted monocyte activity can lead to detrimental hyperinflammation, manifesting as cytokine storms, tissue damage, and even death. Regulating monocyte activity is, therefore, essential for maintaining immune homeostasis, with platelets playing a notable role.

In the blood, platelets surround monocytes, with whom they engage in complex interactions, forming MPAs. These aggregates are common in homeostasis but intensify during inflammation, serving as indicators of inflammatory diseases (Allen *et al*., 2019; Barrett *et al*, 2019; Manne *et al*., 2020). Despite their prevalence, the specific roles of MPAs in both health and disease remain to be fully elucidated.

Our research has uncovered that platelets are integral to sustaining the inflammatory responses of monocytes. Depletion of platelets from human monocytes results in an anergic state, with reduced cytokine production and downregulation of pro-inflammatory genes following stimulation of pattern recognition receptors (PRRs). This hyporesponsive state is also observed in monocytes from ITP patients, who have an impaired ability to produce cytokines in response to PRR activation. Remarkably, reintroducing autologous platelets to platelet-depleted monocytes reversed monocyte immunoparalysis, underscoring the crucial role of platelets in modulating monocyte function. We extended these investigations to the clinical domain by demonstrating the ability of healthy platelets to revert the immunoparalysis of ITP monocytes.

Immunoparalysis is commonly observed in occasions of increased blood loss, such as trauma, major organ surgery (Haupt *et al*, 1998), hemorrhagic fevers (Vangeti *et al*, 2021), sepsis (Cao Yu & Chai, 2019; Weisheit *et al*, 2020), liver failure (Lin *et al*, 2007; Wasmuth *et al*, 2005), and more recently in SARS-CoV-2 infections (Agrati *et al*., 2020; Arunachalam *et al*., 2020). Thrombocytopenia is a common feature among these disparate conditions. However, additional confounding factors, such as co-infections and lower leukocyte counts, render it difficult to assess the actual contribution of platelets to monocyte immunoparalysis. Our study provided evidence of monocyte immunoparalysis in ITP patients without other confounding factors such as infections or glucocorticoids and with average leukocyte counts and C-reactive protein levels.

Our findings reinforce the longstanding belief that normal platelet counts are essential for maintaining immune homeostasis. Low platelet count is a significant risk factor for infections (Birnie *et al*, 2019; de Stoppelaar *et al*., 2014), and ITP patients experience compromised host defense, which is associated with a higher risk of infection, poorer treatment response, and more extended hospital stays (Qu *et al*, 2018). In spite of that, systemic levels of IL-1β, IL-6, and TNFα in ITP patients are usually comparable to healthy individuals (Andreescu, 2023). Our findings *in vivo* recapitulate the human scenario and suggest that platelet depletion skews the immune response, prioritizing cell survival over host defense. Platelet-depleted mice exhibited no significant differences in plasma levels of pro-inflammatory cytokines or in their monocyte transcriptional response to intraperitoneal LPS. However, monocytes from these mice displayed weakened transcription programs associated with host defense. Although contradictory to the transcriptional profile of platelet-depleted human monocytes, our findings *in vivo* are fully corroborated by mouse models of sepsis, where thrombocytopenia is associated with disturbed immune defense programs and increased mortality independently of disease severity but without influencing local or systemic inflammation, plasma cytokine levels, and neutrophil recruitment (Carestia *et al*., 2019; Claushuis *et al*., 2016; van den Boogaard *et al*., 2015; Xiang *et al*., 2013). Furthermore, and in agreement with our findings, platelet depletion resulted in enhanced coagulation, endothelial cell activation, and even increased plasma levels of some pro-inflammatory cytokines (Carestia *et al*., 2019; Claushuis *et al*., 2016; van den Boogaard *et al*., 2015; Xiang *et al*., 2013). Overall, our findings highlight platelets as essential checkpoints determining the magnitude of monocyte innate immune programs and may have consequences for the treatment of immunosuppression or other life-threatening conditions associated with unrestrained monocyte responses (Ferreira et al., 2021; Junqueira et al., 2022).

The interaction between platelets and monocytes enhances various monocyte functions, such as their ability to migrate through vessel walls, produce reactive oxygen species, and differentiate into dendritic cells, ultimately aiding in the activation of CD8+ T cells (D’Mello *et al*., 2017; Fu *et al*., 2021; Han *et al*., 2020; Rong *et al*., 2014; Singhal *et al*., 2017). However, the molecular mechanisms of how platelets influence these monocyte functions remain partially understood (Kral *et al*, 2016). Our study introduces a novel mechanism of platelet regulation on monocyte immunity, highlighting platelet-derived NFκB as a key trans-cellular regulator of cytokine production in monocytes. Adding fresh platelets counteracted genetic ablation of NFκB signaling in human monocytes. Furthermore, while pharmacological NFκB inhibition was toxic to primary human monocytes, we found that platelets bypass SB203580 inhibition of p38 MAPK. These findings align with existing literature that specific ablation of p38 MAPK in platelets confers protection against myocardial infarction (Shi *et al*., 2017) and platelet-activating factor (PAF)-induced lethality (Abhilasha *et al*, 2019). Indeed, MAPK p38 inhibitors have emerged as pertinent targets for cytokine-suppressive anti-inflammatory drugs (Anand *et al*, 2011; Christie *et al*, 2015; Yong Koh & Moon, 2009).

We also corroborated previous findings indicating a rich stock of NFκB subunits in platelets and MKs (Beaulieu & Freedman, 2009; Lannan *et al*., 2015; Spinelli *et al*, 2010). Transcriptional and kinome analyses have provided insight into the molecular underpinnings of this interplay, revealing that platelets contribute to key cytokine-regulating signaling pathways in monocytes, notably NFκB and MAPK. Interestingly, it has been previously demonstrated that interaction with platelets enhances the nuclear translocation of p65 within monocytes (Ueda *et al*, 1994; Weyrich *et al*, 1996; Weyrich *et al*., 1995). However, the possibility that a considerable portion of p65 arises from platelets was neither raised nor experimentally demonstrated. Our study provides the first evidence for the translocation of NFκB from platelets to monocytes, with maintained functions in monocytes. We have shown that platelets can reactivate TLR-induced cytokine secretion in RelA^−/−^ and NFκB2^−/−^ THP1 monocytes, pinpointing platelets as the only viable source of these molecules.

Although platelets could also reconstitute cytokines in SB203580-treated monocytes, our mass speck proteomics approach did not identify platelet-derived p38a MAPK on human monocytes. However, as MS is an abundance-based method, we cannot exclude the transfer of low anounts of p38a contributing to pathway activity. As p38 MAPK signaling is required for the transcriptional NFκB activity (Anderson, 2010; Gottschalk *et al*, 2016; Guma *et al*, 2011; Saha Jana & Pahan, 2007), platelets may circumvent SB203580 inhibition in monocytes by supplying NFκB. Supporting this hypothesis, SB203580 reduced the phosphorylation of p65 (RelA) in treated monocytes (Fig. **S7E**).

Although the transfer of TFs from platelets to monocytes has been anticipated (Lannan *et al*., 2015), it has never been experimentally demonstrated. In this regard, our study establishes platelet-monocyte TF transfer as a sophisticated mechanism of intercellular communication. This finding is particularly intriguing given the unique presence of platelets in mammals, as opposed to invertebrates, which rely on a single-cell type for host defense and wound healing.

Although we ruled out the participation of numerous well-known ligand-receptor interactions in the platelet-monocyte crosstalk, the mechanisms by which platelets supply TFs to monocytes remain unidentified. Though we have not visualized phagocytosis of platelets by monocytes by confocal microscopy, using phagocytosis inhibitors, and transfer of platelet supernatants, we ruled out the involvement of a humoral platelet-derived factor, supporting that vesicles restore cytokine activity in platelet-depleted monocytes, a known mechanism for the transfer of cellular receptors (Rozmyslowicz *et al*, 2003), bioactive lipids (Barry *et al*, 1997; Rossaint *et al*, 2016) and intact organelles (Levoux *et al*, 2021) that are transferred from platelets, or platelet vesicles, to leukocytes, with preserved activity in the cytosol of recipient cells. We also observed numerous platelets bound to monocytes, in line with previous observations that a significant fraction of blood monocytes (up to 45%) are physiologically associated with platelets in steady-state conditions (Rinder *et al*., 1991). Furthermore, breaking MPAs via interfering with P-sel-PSGL-1 interactions did not prevent the platelet effect on monocytes. However, based on our experiments, we cannot exclude that cell-surface receptor-activated signaling cascades involving NFκB contribute to platelet-induced monocyte activation. In a randomized clinical trial, two anti-platelet medications reduced MPA formation and the systemic levels of CCL2, IL-6, IL-8, and TNFα in humans challenged with intravenous LPS injection (Thomas *et al*, 2015). However, depending on the disease setting, platelets attenuate monocyte inflammatory responses by triggering the production of IL-10 or sequestering IL-6 and TNF-α released by monocytes (Carestia *et al*., 2019; Xiang *et al*., 2013). Hence, platelet-monocyte interactions can have different and context-dependent outcomes (Kral *et al*., 2016). Our findings further indicate that other classical and well-described mechanisms of platelet-monocyte crosstalk (i.e., co-stimulatory molecules, integrins, CD40-CD40L axis, CCL5, CXCL12, and SAs) are not involved in the effects described. The exclusion of these mechanisms highlights a previously unrecognized mechanism of immune regulation, where platelets license the pro-inflammatory cytokine responses of human monocytes. Our findings indicate that the complex association between monocytes and platelets may be part of a homeostatic mechanism to equip monocyte immune functions when needed.

In summary, our study offers a novel perspective on the regulatory role of platelets in monocyte-mediated immunity. By identifying platelet-derived NFκB and MAPK signaling as crucial for monocyte cytokine production, we open the door to potential therapeutic strategies that could modulate this platelet-monocyte interaction to manage inflammatory conditions and improve outcomes in diseases characterized by monocyte dysfunction. Our findings may also help to pave the way for new clinical approaches for improving platelet transfusions, which are usually safe but can occasionally have deleterious inflammatory consequences.

## METHODS

### Reagents

Cell culture reagents (e.g., PBS, Fetal calf serum, GlutaMax, and RPMI) were from Gibco, Thermo Fisher Scientific. Stimuli were as follows: LPS (Ultrapure from E.coli O111:B4), Pam4CysK4, and R848 (Resiquimod) were from Invivogen. Nigericin acid was from Thermo Fisher Scientific. Inhibitors used were Cytochalasin D, BAY 11-7082 (Sigma-Aldrich). The CellTiter-BlueTM Cell Viability Assay and Caspase-Glo. 1 Inflammasome Assay was from Promega. HTRF kits to detect IL-1b, IL-6, TNFa, and NFκB were from Cisbio. ProcartaPlexTM kits for detecting other Cytokine/Chemokine/Growth Factors were from Invitrogen. Antibodies were anti-IL-1b antibody (BAF401, R&D Systems, 1:1,000). DRAQ5 was from Thermo Fisher Scientific. For FACS, we used FcR Blocking Reagent, human and mouse (Miltenyi Biotec). Anti-human CD41 directly conjugated to Alexa Fluor 488 (Clone: HIP8), Anti-human CD14-A647 (Clone: HCD14), and isotype Mouse IgG1, k Isotype Ctrl (Clone: MOPC-21) Antibody were all from BioLegend.

### Study Subjects

Peripheral blood was obtained by venipuncture of healthy volunteers after the signature of informed consent and approval of the study by the Ethics Committee of the University of Bonn (Protocol #282/17), following the Declaration of Helsinki. Patients with immuno-thrombocytopenia were recruited from a tertiary hematology referral hospital. We recruited five patients (3 females and two males, average age of 46 ± 9) from December 2019 to May 2020. All patients were diagnosed with ITP, and comorbidities/infections were ruled out. Written informed consent was obtained from all subjects according to the Declaration of Helsinki and approval by the Institutional Review Board of the University of Bonn (313/21). All participants were instructed about the study’s objectives and signed an informed consent following guidelines for human research.

### Primary cells

Primary human monocytes were isolated from peripheral blood collected from healthy volunteers in S-Monovette  K3EDTA tubes. Blood was diluted 1:3 in PBS, and peripheral blood mononuclear cells (PBMCs) were isolated using Ficoll^®^ Paque PLUS density gradient centrifugation at 700 x g, 20 min. Primary human CD14+ monocytes were isolated from PBMCs using positive magnetic separation with the Classical Monocyte Isolation Kit (Miltenyi Biotech) or negative selection with the EasySep™ Human Monocyte Isolation Kit (STEMCELL^TM^ Technologies) following manufacturer instructions. Platelet-depleted monocytes were generated by adding a Platelet Removal Component (50 µl ml^−1^) supplied with the isolation kit (STEMCELL^TM^ Technologies). The purity of monocyte populations was assessed by flow cytometry. Cells were incubated with FcR Blocking Reagent for 15 min at 37 °C, followed by staining with fluorochrome-conjugated monoclonal antibodies against human CD14 APC (eBioscience), anti-CD45 PE (eBioscience), and anti-CD41a FITC (eBioscience) for 30 min at RT. Cells were washed with PBS, and fluorescence was measured on a MACSQuant® Analyzer 10.

### Platelet isolation

Human platelets were isolated from venous blood from healthy volunteers collected as previously described**^(22, 87)^**. Briefly, whole blood was centrifuged at 330 x g for 15 min, and platelet-rich plasma (PRP) was collected. To prevent platelet aggregation during isolation, PRP was treated with 200 nM Prostaglandin E1 (PGE1), diluted 1:1 with PBS, and centrifuged at 240 x g for 10 min to pellet leukocytes. Platelet suspensions were collected and washed by centrifugation at 430 xg for 15 min in the presence of PBS PGE1. Washed platelets were pellet and resuspended in pre-warmed RPMI. Human platelets were stimulated with 1 U ml−1 Thrombin from human plasma (Sigma-Aldrich) or left unstimulated for 30 min to assess the purity and activation of isolated cells. Platelets were incubated with FcR Blocking Reagent followed by staining with CD45 (leukocyte marker), and CD41a FITC (eBioscience) and anti-CD62P APC (eBioscience).

### Generation of platelet supernatants

Platelet supernatants were generated from suspensions of 1 x 10^8^ human platelets incubated in RPMI and left untreated (PltSups) or stimulated with 2 ng ml−1 LPS (PltsLPS-Sups) for 4.5 h at 37°C. Cell suspensions were centrifuged at 3,000 x g for 10 min. The platelet supernatants were harvested and used to stimulate human monocytes.

### Cell lines

The human monocytic THP-1 cell line (ATCC TIB-202) and the THP-1 DualTM reporter cell (InvivoGen, Thpd-nfis) were culture in RPMI supplemented with 10% heat-inactivated fetal calf serum, 1% Penicillin/Streptomycin and 1x GlutaMax. For THP-1 DualTM cells, the growth medium contained additionally 25 mM HEPES. Cells were either studied in a monocyte-like phenotype (without PMA-differentiation) or after their differentiation into macrophages by PMA (50 µM) treatment for 24 h.

### Stimulation Assays

Monocytes (1 x 10^5^/well) were cultured alone or co-cultured with 50 (5 x 10^6^/well) to 100 platelets (1 x 10^7^/well). For stimulation, cells were treated with agonists of TLR1/2: Pam3CysK4 (1 µg ml^−1^); TLR4: LPS (2 ng ml^−1^); or TLR7/8: Resiquimod R848 (3.5 µg ml^−1^); for for 4.5h. For inflammasome activation, cells were primed with a TLR agonist for 3 h (indicated in Figs), followed by activation with nigericin (10 µM, 90 min) PrgI from *Bacillus anthracis* (100 ng ml^−1^) and protective antigen (PA, 1 µg ml^−1^), for 90 min) or transfection of poly(dA:dT, 0.5 µg ml^−1^, 3 h). After stimulation, CFS was harvested and used for cytokine detection.

### Cytokine measurement

According to the manufacturer’s instructions, cytokines were detected in cell-free supernatants or whole cell lysates by homogeneous time-resolved fluorescence technology (HTRF, Cisbio). A Human ProcartaPlexTM (Invitrogen) immunoassay was additionally used to detect 45 human cytokines, chemokines, and growth factors, according to the manufacturer’s instructions, using a MAGPIX® System.

### Cell Viability Assays

Cell viability was determined by measuring the release of Lactate Dehydrogenase Assay (LDH) in CFS or the CellTiter-Blue Assay (CTB) in stimulated cells. For LDH measurement, CFS from stimulated cells were incubated with LDH buffer for 30 min at 37 °C without CO2, and absorbance was measured at 490 nm using a Spectramax i3 plate reader (ThermoFisher). Cells were incubated with CTB buffer at 37°C, and the fluorescence was read at different time points in both assays, cells treated with Triton X-100 served as a 100% cell death control. Fluorescence was detected with a SpectraMax.

### Assessment of Caspase-1 Activity

The specific activity of Caspase-1 was assessed in cell-free supernatants using Caspase-Glo® 1 Inflammasome Assay (Promega). Briefly, CFS was mixed 1:1 with Glo Buffer and incubated for 30 min at RT. Luminescence was measured with SpectraMax i3 (Molecular devices).

### Confocal Imaging

Platelet interactions with human monocytes were imaged in a Leica TCS SP5 SMD confocal system (Leica Microsystem). Cells were seeded in microslide 8-Well IBIDI chamber, fixed with 4% paraformaldehyde (PFA) for 30 min, washed and incubated with staining mAbs in permeabilization/blocking buffer (10% goat serum, 1% FBS, and 0.5% Triton X-100 in PBS) supplemented with human FcR Blocking Reagent for 30 min. Cells were stained overnight at 4°C with anti-human CD41Alexa Fluor 488, anti-human CD14-A647, or Hoechst 34580 (ThermoFisher).

### Immunoblotting

Cells were lysed with RIPA complete lysis buffer supplemented with EDTA-free protease and phosphatase inhibitors (Roche) and 25 U of Benzonase Nuclease. Samples were loaded onto NuPAGETM Novex 4-12% Bis-Tris gels and transferred to Immobilon-FL Polyvinylidene fluoride (PVDF) membranes. After blocking with 3% BSA [w/v] in Tris-buffered saline (TBS), membranes were incubated overnight with primary antibodies diluted in TBS containing 3% BSA [w/v] and 0.1% Tween. After washing steps, membranes were incubated with secondary antibodies conjugated to IRDye680 or IRDye800 (1:25000) for 2 h at RT. The membrane was scanned at Odyssey Imager (Li-Cor Biosciences). The following primary antibodies were used for: β-Actin rabbit mAb (1:5000), Phospho-NFκB p65 (Ser536) (93H1) Rabbit mAb #3033 (1:1000), NFκB p65 (C22B4) Rabbit mAb #4764 (1:1000) and IkBa (L35A5) Mouse mAb (Amino-terminal Antigen) #4814 (1:1000).

### Platelet depletion *in vivo*

In vivo, platelet depletion was performed in female C57BL/6J mice (12 weeks) by intravenousl (i.v.) injection of 2 mg/kg of polyclonal rat anti-mouse GPIbα (R300) or same amounts of a polyclonal rat IgG none-immune (C301) antibody as control (Emfret Analytics). Mice were then challenged 12 hours later with i.v. injection of 2 mg/kg LPS or PBS as control. Blood was collected after 3 h, and PBMCs were isolated by density gradient centrifugation in Ficoll Paque PLUS. Mouse monocytes were separated from PBMCs by FACS-sorting gating on Ly6G^−^, IA/IE^−^, CD45^+^, CD11b^+^, and CD115^+^ cells and sorted directly into QIAzol lysis reagent. Lysates were frozen in liquid nitrogen and sent to GENEWIZ Azenta Life Sciences for bulk RNA sequencing. All animal experimentation was approved by the local ethical committee (LANUV-NRW #84-02.04.2016.A487). Mice were purchased from Charles River and housed at the House for Experimental Therapy (HET) of the University Hospital of Bonn.

### Fluorescence-activated cell sorting was used to obtain mouse monocytes

Mouse blood monocytes were isolated from pooled blood of 4 mice per experimental group in 3 different experiments. For this, mouse PBMCs were isolated through Ficoll gradient and stained with anti-Ly6G BV785 (Biolegend, dilution: 1:40), anti-I-A/I-E BV785 (Biolegend, dilution: 1:80), anti CD45 FITC (Biolegend, dilution: 1:200), anti-CD11 b PE (eBioscience, dilution: 1:166), anti-CD115 APC (Biolegend, dilution: 1:80) and Hoechst 33258 (Abcam, 1 µM) as viability staining. Finally, viable Ly6G-I-A/I-ECD45+ CD11b+CD115+ monocytes were sorted directly into QIAzol lysis reagent using the BD FACS Aria Fusion (387333827) and BD FACS Aria III (216372545) at the Flow Cytometry Core Facility of the Medical Faculty of the University of Bonn.

### Bulk RNA sequencing of isolated murine monocytes

For bulk RNA sequencing, samples were processed according to GENEWIZ (Azenta Life Sciences) pipeline. RNA extraction was performed, followed by a library preparation. Ultra-low input RNA-Seq was performed at Illumina HiSeq PE 2x150 bp with ∼350M reads. Reads were then mapped to the Mus musculus GRCm38 reference genome (ENSEMBL) using the STAR aligner v.2.5.2b, and unique gene hit counts were generated with the featureCounts from the Subread package v.1.5.2. Finally, standard analysis was performed by GENEWIZ. Differential gene expression analysis was performed using DESeq2, p-values, and log2 fold changes were calculated by applying the Wald test, and genes with adjusted p-value < 0.05 and absolute fold change > 2 were considered significantly altered differentially expressed genes (DEGs). Gene ontology (GO) analysis of the significant DEGs was applied by GENEWIZ (Azenta Life Sciences) using the Fisher exact test. GO terms were assigned using the GeneSCF v.1-p2 software. Data visualization was created using the R package ggplot2.

### Transcriptional profiling of human monocytes and platelets with NanoString

According to the manufacturer protocol, the nCounter® Human Myeloid Innate Immunity Panel v2 (NanoString® Technologies) was used to assess the mRNA expression of 770 human transcripts. Briefly, single or co-cultures of monocytes and platelets were lysed in RLT Buffer (Qiagen) containing β-Meracptoethanol (1 x 10^4^ cells/µl). RNA was isolated with and homogenized with CodeSets and left for hybridization overnight. RNA/CodeSet complexes were immobilized on nCounter cartridges at the nCounter® Prep Station 5s, and data collection (RCC files) and quality check was performed in the nCounter® Digital Analyzer 5s. The nSolver^TM^ Analysis and Partek® Genomics Suite® software was used for analysis.

### Kinase activity profiling microarray

Primary monocytes and platelets were isolated from 6 healthy donors. Monocytes were stimulated *ex vitro* before or after the removal of platelets or after the re-addition of platelets to PdMo (biological replicates with n = 6). As before, platelets alone (n = 6) were assessed. Stimulated cultures were lysed in M-PER mammalian protein extraction reagent supplemented with PhosphoSTOP (Roche) and cOmplete Tablets (Roche) incubated on ice for 15 minutes. After centrifugation at 20.000x g for 15 minutes at 4 C, the protein concentration in the supernatant is evaluated via Bradford assay. The lysate is aliquoted, snap-frozen, and stored at −80 C. Only unthawed aliquots are used for the kinase activity assay.

Ser/Thr Kinase (STK) activity profiling assays based on measuring peptide phosphorylation by protein kinases (PamGene International BV, The Netherlands) were performed as instructed by the manufacturer. In summary, samples with 1 μg protein were applied on PamChip 4 arrays containing 144 (STK) or 196 (PTK) peptides immobilized on a porous aluminum oxide membrane. The peptide sequences (13 amino acids long) harbor phosphorylation sites and are correlated with one or multiple upstream kinases. Fluorescently labeled anti-phospho antibodies detect phosphorylation activity of kinases present in the sample. Instrument operation and imaging are controlled by the EVOLVE 2.0 software and quantified using BioNavigator 6.3 (BN6; PamGene International BV, The Netherlands). Signal intensities at multiple exposure times were integrated by linear regression (S100), Log2-transformed, and normalized using a Combat correction model for batch correction where the scaling parameters (mean / sd) are estimated using an empirical Bayes approach, which effectively reduces the noise associated with applying the correction.

The normalized values were used to perform statistics comparing groups or the upstream kinase analysis (UKA) tool (BN6; PamGene International BV). The following statistical test is used to generate a list of differentially significant phosphorylated peptides: Paired T-test for LPS effect/ comparisons and ANOVA-PostHoc Test for multiple treatments versus control for comparing each group to StdMo unstimulated in both unstimulated and LPS-15 min conditions. For Kinase interpretation (differential): To generate a ranked list of putative kinases responsible for differences in the peptide phosphorylation, PamGene’s in-house method called Upstream Kinase Analysis (UKA) was used. For Pathway interpretation (differential): To generate a ranked list of possible canonical pathways (and networks) responsible for differences in the peptide phosphorylation, there are many open-source tools to perform this.

The phylogenetic kinome tree, applicable to group the kinases into sequence families, is plotted using the online portal CORAL: http://phanstiel-lab.med.unc.edu/CORAL/ (12). The upstream kinase analysis functional scoring tool (PamGene International) rank-orders the top kinases differential between the two groups, the ranking factor being the final (median) kinase score (represented by node size). This score is based on a combined sensitivity score (difference between treatment and control groups, expressed as node color) and specificity score for a set of peptides to kinase relationships derived from existing databases. An arbitrary threshold of a final score 1.2 was applied based on the specificity scores. Significant peptides (t-tests, p-value < 0.05) or kinases (UKA, final scores > 1.2) were imported to the MetaCore pathway analysis tool (Clarivate Analytics), where an enrichment analysis was performed for pathways and networks. It matches the kinases or substrates in the kinome array data with functional ontologies in MetaCore. The probability of a random intersection between a set of IDs in the target list with ontology entities is estimated in the p value of hypergeometric intersection. The lower p-value means higher relevance of the entity to the dataset, which shows a higher rating for the entity. Direct interaction network algorithms were used to build interconnected networks within each comparison. The “Add interactions” feature was used to add the interaction between RIPK1 and the data in the MetaCore™ database after it was built.

### Statistical Analysis

Statistical analyses were performed with GraphPad Prism Version 9.0f. Unless indicated otherwise, all graphs are built from pooled data from a minimum of two independent experiments (biological replicates) performed in triplicates (technical replicates). All statistical analyses were preceded by normality and lognormality tests, followed by the recommended parametric or nonparametric tests. For most experiments with several groups (StdMo, PdMo, PdMo + Ptls, and Plts, stimulated vs. unstimulated), P values were determined by two-way ANOVA with Tukey’s or Sidak’s or multiple comparison tests. No outliers were detected or removed from the analysis. Additional statistical details are given in the respective Fig legends when appropriate.

### Data sharing availability

All datasets that support this manuscript will be made fully available upon publication. Raw data for RNA-seq datasets are being deposited into public databases, and we will share with reviewers the assessment number upon request. Additional requests to assess raw data and transcriptomic datasets before publication should be made via e-mail to the corresponding author, along with an analysis proposal.

### Data Presentation

Unless indicated otherwise (in Fig legends), all graphs are represented as Floating Bars (with mean and minimum to maximum values) and are built from pooled data from a minimum of two independent experiments (biological replicates) performed in triplicates (technical replicates) with monocytes or platelets from different donors. Each symbol represents the average from 3 technical replicates per donor or experiment in the case of cell lines. Symbols are coded (●, ▾, ◼ □□, ●, etc.) to indicate donors so readers can track the internal variability between different donors or experiments. Dots are semi-transparent, with darker symbols showing overlapping points.

## FUNDING

This study was funded by the European Research Council (EC | H2020 | H2020 Priority Excellent Science | H2020 European Research Council (ERC) PLAT-IL-1, 714175). Bernardo S Franklin is further supported by the HORIZON Lump Sum Grant (ERC PoC UNBIAS, 101123144), (BEATSep, 101137484), as well as Germany’s Excellence Strategy (EXC 2151 – 390873048, SFB TRR259) from the Deutsche Forschungsgemeinschaft (DFG, German Research Foundation).

## ACKNOWLEDGMENTS

We thank Marco A. Ataide, Florian I Schmidt, and Dagmar Wachten for scientific discussions and input and for proofreading the manuscript. We thank Cornelia Rohland and Maximilian Rothe for technical help and organization and maintenance of mouse lines, respectively. We thank Jonathan Schmid-Burgk for their advice and support in generating THP1 KO cells using CRISPR-Cas9. We thank Dr. Savithri Rangarajan and Dr. Rik de Wijn (PamGene, Diagnostic Assay Services, ’s-Hertogenbosch, The Netherlands) for performing the peptide quality control and bioinformatics analysis of the Protein Kinase Assay. We thank the Microscopy and Flow Cytometry Core Facility of the Medical Faculty at the University of Bonn for providing help, services, and devices funded by the Deutsche Forschungsgemeinschaft (DFG, German Research Foundation, project number 388158066, 216372545 and 388159768). Finally, we thank Andrea Schlichting and Leonie Verwohlt for administrative support.

## AUTHOR CONTRIBUTIONS

**Ibrahim Hawwari**: Investigation; Formal analysis; Data curation; Visualization; Methodology; Writing—review and editing. **Lukas Rosnagel:** Investigation; Formal analysis; Data curation; Visualization; Methodology; Writing—review and editing. **Nathalia Sofia Rosero Reyes:** Investigation; Technical Support. **Lino L Teichmann** and **Lisa Meffert**: ITP patient samples. **Agnieszka Demczuk:** Investigation. **Lucas S. Ribeiro:** Technical Support. **Damien Bertheloot:** Supervision; Investigation. **Magali Noval Rivas**: Investigation. **Moshe Arditi:** Formal analysis; Data curation. **Sebastian Kallabis:** Investigation; Formal analysis; Data curation. **Felix Meissner:** Formal analysis; Data curation. **Bernardo S Franklin:** Conceptualization; Funding acquisition; Project administration; Resources; Formal analysis; Supervision; Validation; Investigation; Visualization; Writing—original draft.

## DISCLOSURE AND COMPETING INTERESTS STATEMENT

All authors declare that they have no conflict of interest.

Graphs display floating bars with the max/min values and mean (white bands) combined from multiple experiments. P values were calculated with 2-Way Anova, Tukey’s multiple comparison test, and are displayed in the Fig. Each symbol represents one independent experiment or blood donor.

## Notes

### Competing Interest Statement

The authors have declared no competing interest.

### Summary of Updates

Figure 4 was missing in the submitted manuscript.

