## Supporting Figures for "Platelet-derived transcription factors license human monocyte inflammation"

#### Dataset 1

**Blocking strategies to assess the involvement of immune co-stimulatory molecules, Integrins, CD40-CD40L, and Sialic Acids in the platelet-effect.**

Platelets produce RANTES (CCL5) and SDF1a (CXCL12), which regulate inflammatory functions of monocytes(Alard et al, 2015; Chatterjee et al, 2015). However, in contrast to platelets, the supplementation of PdMo with rhCCL5 or rhCXCL12 did not rescue their faulty cytokine secretion (Fig S5C). Furthermore, these cytokines were not consistently detected in platelets by our Luminex assays (Figure 1G; Fig S1G).

As demonstrated in our protein kinase assay, the activation of the CD40-CD40L axis figured in the top pathways regulated by platelet-monocyte interactions (Fig 6E). This is consistent with the description of CD40L expression on platelets. CD40L binding to CD40 is a co-stimulatory signal for cytokine secretion in macrophages(Henn et al; Inwald et al, 2003). However, supplementation of PdMo with soluble rhCD40L did not substitute platelets in their ability to rescue the impaired cytokine secretion (Fig S5A). Furthermore, blockage of the CD40-CD40L axis with anti-CD40L mAb had no influence on the platelet rescue of cytokines in PdMos (Fig S 5B-C). These findings conclusively exclude the CD40-CD40L axis as the mechanism for the interdependency of monocytes on platelets for their cytokine responses.

Platelets contain sialic acids (SAs), which are implicated in aggregation and adhesion to leukocytes. To examine whether platelet SAs influences monocytes-derived cytokine response, we generated desialylated platelets (dsPlts) by Neuraminidase from *Arthrobacter ureafaciens* treatment. dsPlts were equally able to rescue cytokine secretion in PdMo as intact platelets. Consistent with previous reports of hyperactivity in dsPlts(Kullaya et al, 2018), supplementation of PdMo with dsPlts induced higher cytokine levels as intact platelets (Fig S 5D). To further examine the role of SAs in the platelet-monocyte crosstalk, we tested whether platelets crosslink SA-binding Ig-like lectin-7 (Siglec-7) to drive a pro-inflammatory state of monocytes(Varchetta et al, 2016). However, neither plate-bound nor soluble anti-Siglec-7 rescued the impaired cytokine secretion in PdMos (Fig S 5E).

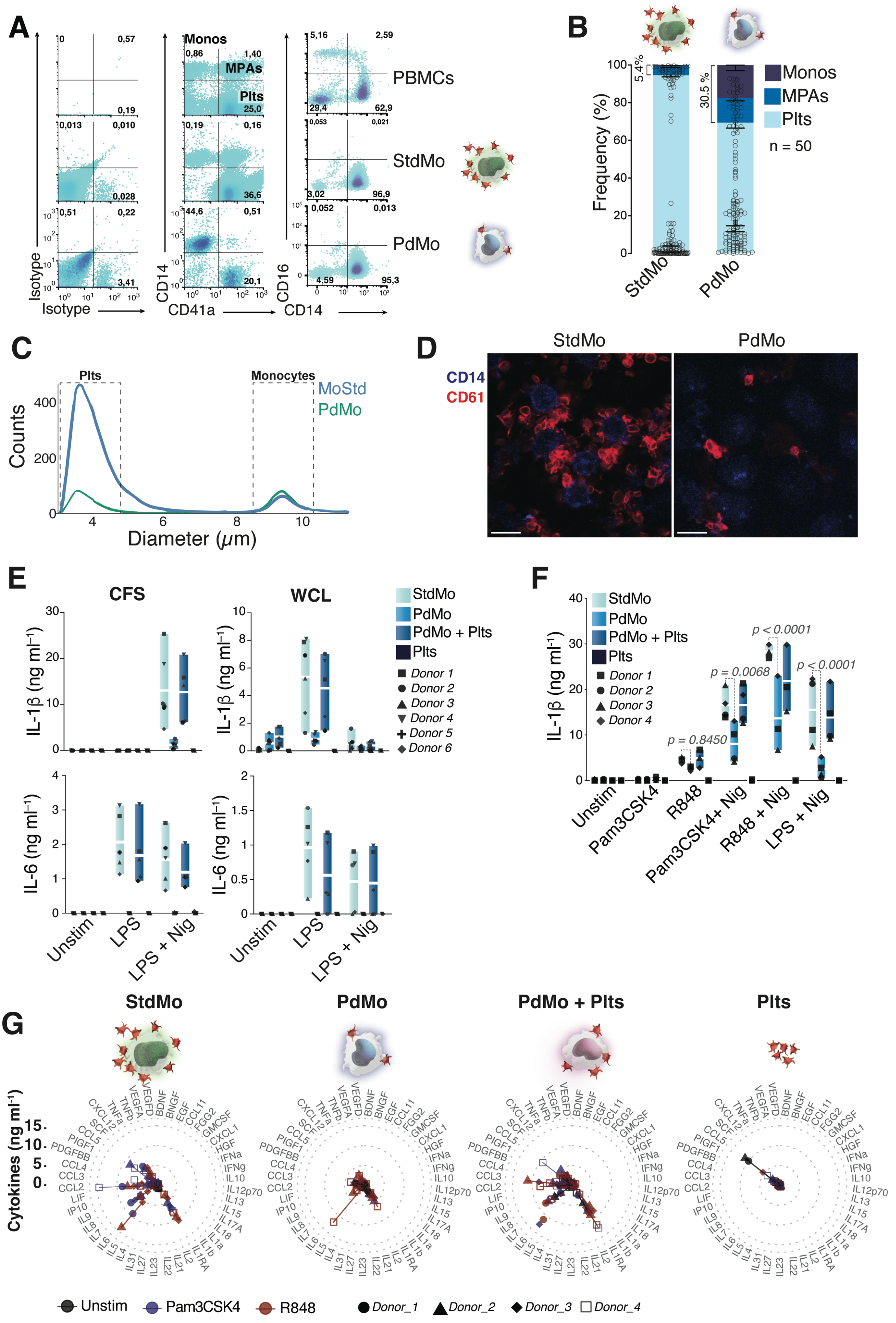
A
Monos
MPAs
Plts
PBMCs
StdMo
PdMo
Isotype
CD14
CD41a
CD16
B
Frequency (%)
5,4%
30,5 %
Monos
MPAs
Plts
n = 50
StdMo
PdMo
C
Plts
Monocytes
MoStd
PdMo
Counts
Diameter (μm)
D
StdMo
PdMo
CD14
CD61
E
CFS
WCL
StdMo
PdMo
PdMo + Plts
Plts
Donor 1
Donor 2
Donor 3
Donor 4
Donor 5
Donor 6
IL-1β (ng ml⁻¹)
IL-6 (ng ml⁻¹)
Unstim
LPS
LPS + Nig
F
StdMo
PdMo
PdMo + Plts
Plts
Donor 1
Donor 2
Donor 3
Donor 4
IL-1β (ng ml⁻¹)
p = 0.8450
p = 0.0068
p < 0.0001
p < 0.0001
Unstim
Pam3CSK4
R848
Pam3CSK4+ Nig
R848 + Nig
LPS + Nig
G
StdMo
PdMo
PdMo + Plts
Plts
Cytokines (ng ml⁻¹)
Unstim
Pam3CSK4
R848
Donor_1
Donor_2
Donor_3
Donor_4

**Fig S1. Platelets are critical checkpoints for the cytokine production of primary human monocytes.**

(**A**) Representative flow cytometry analysis of human PBMCs, and isolated untouched (StdMo), or platelet-depleted (PdMo) primary $CD14^+$ monocytes stained with CD14 (monocytes) and CD41a (platelets), or corresponding isotype controls. Gating strategy to identify platelet-free monocytes ($CD14^+$ $CD41a^–$, Monos), monocyte-platelet aggregates ($CD14^+$ $CD41a^+$, MPAs), and free platelets ($CD14^–$ $CD41a^+$, Ptls). Plots on the right shows immunophenotyping of PBMCs, StdMo and PdMo based on the surface expression of CD14 and CD16. Data is from one representative of several independent experiments.

(**B**) Comparative quantification of Monos, MPAs and Plts populations based on their frequency determined by flow cytometry in monocyte isolations (n = 50) as shown in **A**.

(**C**) Quantification of platelet and monocyte counts through a CASY cell counter and analyzer.

(**D**) Confocal imaging of StdMo showing $CD14^+$ monocytes (blue) and $CD61^+$ platelets (red), and the formation of MPAs. Scale bars: 10 μm.

(**E**) IL-1β and IL-6 concentrations in cell-free supernatants (CFS, left) or whole cell lysates (WCL, right) from untouched (StdMo), platelet-depleted (PdMo), or PdMo that were supplemented with autologous platelets (100:1 platelet:monocyte ratio). Cytokine levels secreted by platelets alone (Plts) were measured as control. Cells were stimulated with LPS (2 ng $ml^{-1}$ for 3 h) followed by activation with nigericin (10 μM for 1.5 h, for IL-1β), or directly with LPS (2 ng $ml^{-1}$ for 4.5 h, for IL-6). Floating bars display the max/min values with inications of the mean (white bands). Each symbol represents one independent experimenbt/ blood donor.

(**F**) IL-1β concentrations in CFS of LPS-primed StdMo, PdMo, or PdMo + Ptls that were stimulated with Pam3CSK4 (1 μg $ml^{-1}$), Resiquimod R848 (10 μM), or LPS (2 ng $ml^{-1}$) for 4.5 h and left untreated, or further activated with nigericin (10 μM, for 90 min). Floating bars display the max/min values with inications of the mean (white bands). Each symbol represents one independent experimenbt/blood donor.

(**G**) Radar plots displaying all 45 cytokines, chemokines and growth factors measured by Cytokine Luminex in the CFS of StdMo, PdMo, or PdMo + Ptls and platelets stimulated with Pam3CSK4 (1 μg $ml^{-1}$), Resiquimod R848 (10 μM) Protein concentrations are represented by the spread from inner (0 ng $ml^{-1}$) to outer circles (> 15 ng $ml^{-1}$). Colors represent stimuli (Unstim, dark grey; Pam3csk4, blue; and R848, red). Each symbol represents one independent experiment/blood donor (n = 4 donors).

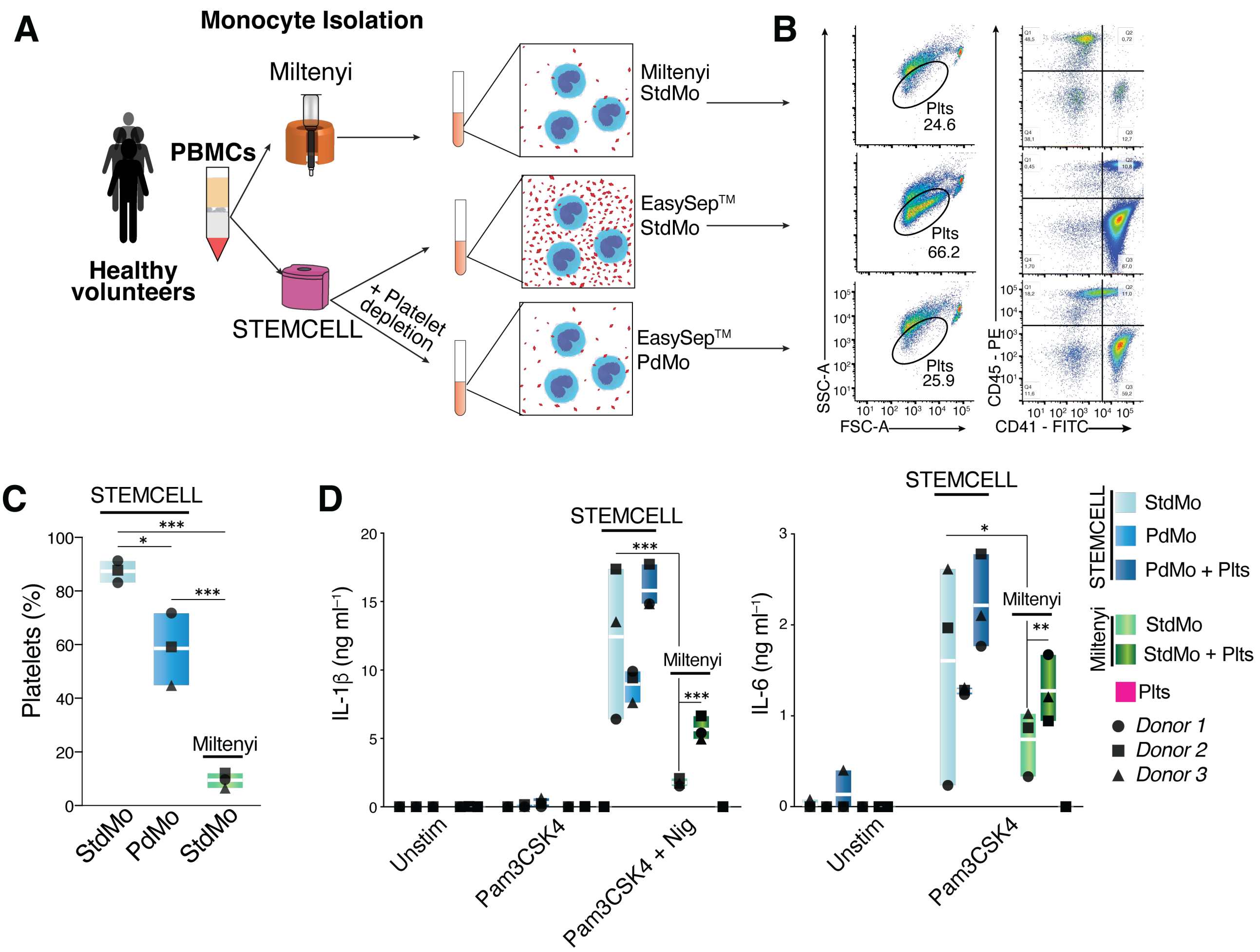


**Fig S2. Impact of platelet removal from primary human monocytes.**

(**A**) Schematic presentation of the immune-magnetic isolation of primary human monocytes from peripheral blood comparing the Miltenyi vs. the EasySep™ monocyte isolation kits. The EasySep™ kit was further supplemented with (PdMo) or without (StdMo) the addition of the platelet-depletion cocktail.

(**B**) Representative flow cytometry analysis of the human primary monocyte populations isolated as in **A**. Gating shows the populations of platelet-free monocytes (CD14$^{+}$ CD41a$^{-}$) and CD41a (platelets), or corresponding isotype controls. Data is from one representative of 3 independent experiments.

(**C**) Frequencies of free platelets in populations of monocytes isolated as in **A**, comparing StdMo (light blue bars), PdMo (blue bars) isolated with the EasySep™ kit, or StdMo isolated with the Miltenyi kit (green bars). Floating bars display the max/min values with indications of the mean (white bands). Each symbol represents one independent experimenbt/blood donor (n = 3).

(**D**) Concentrations of IL-1β, and IL-6 released by untouched (StdMo), platelet-depleted (PdMo), or PdMo that were supplemented with autologous platelets (PdMo + Plts, 100:1 platelet:monocyte ratio), using the EasySep™ kit (blue bars), or untouched isolated with the Miltenyi kit cultured alone (StdMo, geen bars) or co-cultured with platelets (StdMo + Pts). Cytokine levels secreted by platelets alone (Plts) were measured as control. Cells were stimulated with LPS (2 ng ml$^{-1}$ for 4.5 h, for IL-6) or with LPS (3 h) followed by activation with nigericin (10 μM for 1.5 h, for IL-1β). Floating bars display the max/min values with indications of the mean (white bands). Each symbol represents one independent experimenbt/blood donor (n = 3). P values are from Two-way ANOVA with Tukey's multiple comparison test with 95% confidence interval, and are indicated as * (< 0.05), ** (< 0.01), and *** (< 0.001).

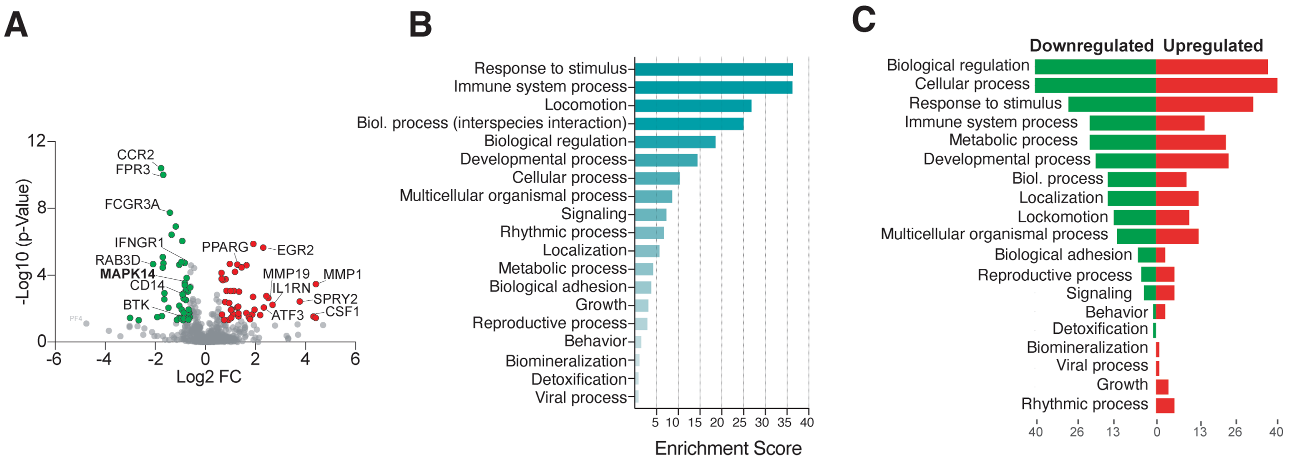


**Fig S3: Platelet depletion induces transcriptional reprogramming in primary human monocytes.**

**(A)** Volcano plot displays differentially expressed genes induced by platelet depletion alone comparing unstimulated MoPD and MoStd. Volcano plot shows significantly upregulated (red) and downregulated (green) genes.

**(B)** Gene ontology (GO) analysis displays enriched gene sets categorized in biological processes that were changed upon platelet depletion.

**(C)** Forest plot presents GO analysis showing the proportions of down- (green) and upregulated (red) enriched gene sets in each GO category. Alterations with fold change ≥ 2 and p-value < 0.05 are considered as significant.

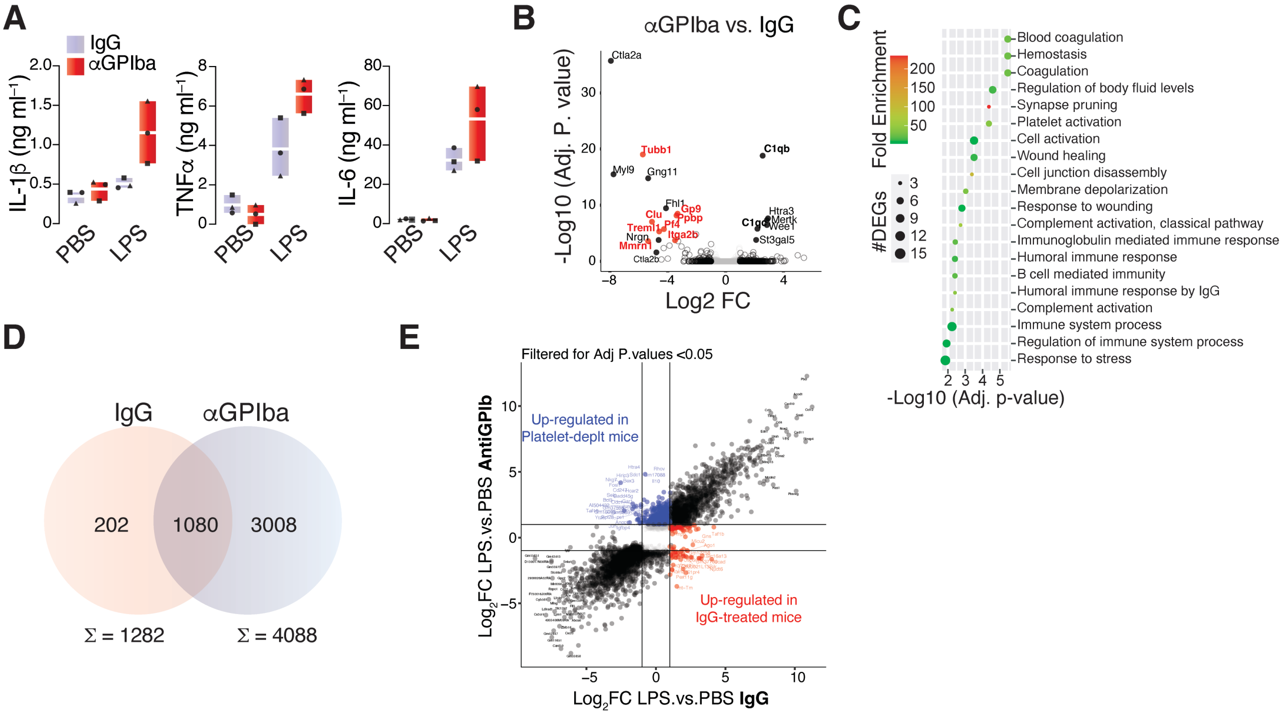


**Fig S4: Bulk RNASeq analysis of FACS-sorted murine monocytes from platelet-depleted mice.**

(**A**) Plasma cytokine levels in IgG or antiGPIbα-treated mice challenged with LPS, or vehicle (PBS).

(**B - C**) Volcano plot showing the log2 fold change (x-axis) and significance (−log10 *p-value; y-axis), and (**C**) pathway enrichment analysis of the differentially expressed genes (DEGs, adjusted P value < 0.05, Fold Change >=2, <= -2) comparing PBS-challenged anti-GPIba-treated vs. IgG-treated mice. (**D**) Platelet-specific transcripts are highlighted in red.

(**D**) Venn-diagram of the DEGs in monocytes taken in IgG (1282) vs. anti-GPIba-treated mice (4088).

(**E**) Volcano plots (log2 fold change vs. −log10 *adjusted P-value) of the significant DEGs (adjusted P-value < 0.05, Fold Change ≤ -2 or ≥ 2) comparing PBS vs. LPS-challenge in IgG (left) or anti-GPIba-treated mice (right).

(**F**) Scatter plot comparing the DEGs (-2 <= Fold Change >= 2, FDR corrected P.value < 0.05) of LPS-induced gene expression in monocytes isolated from IgG-treated (x.axis) vs. anti-GPIbα-treated mice (y.axis).

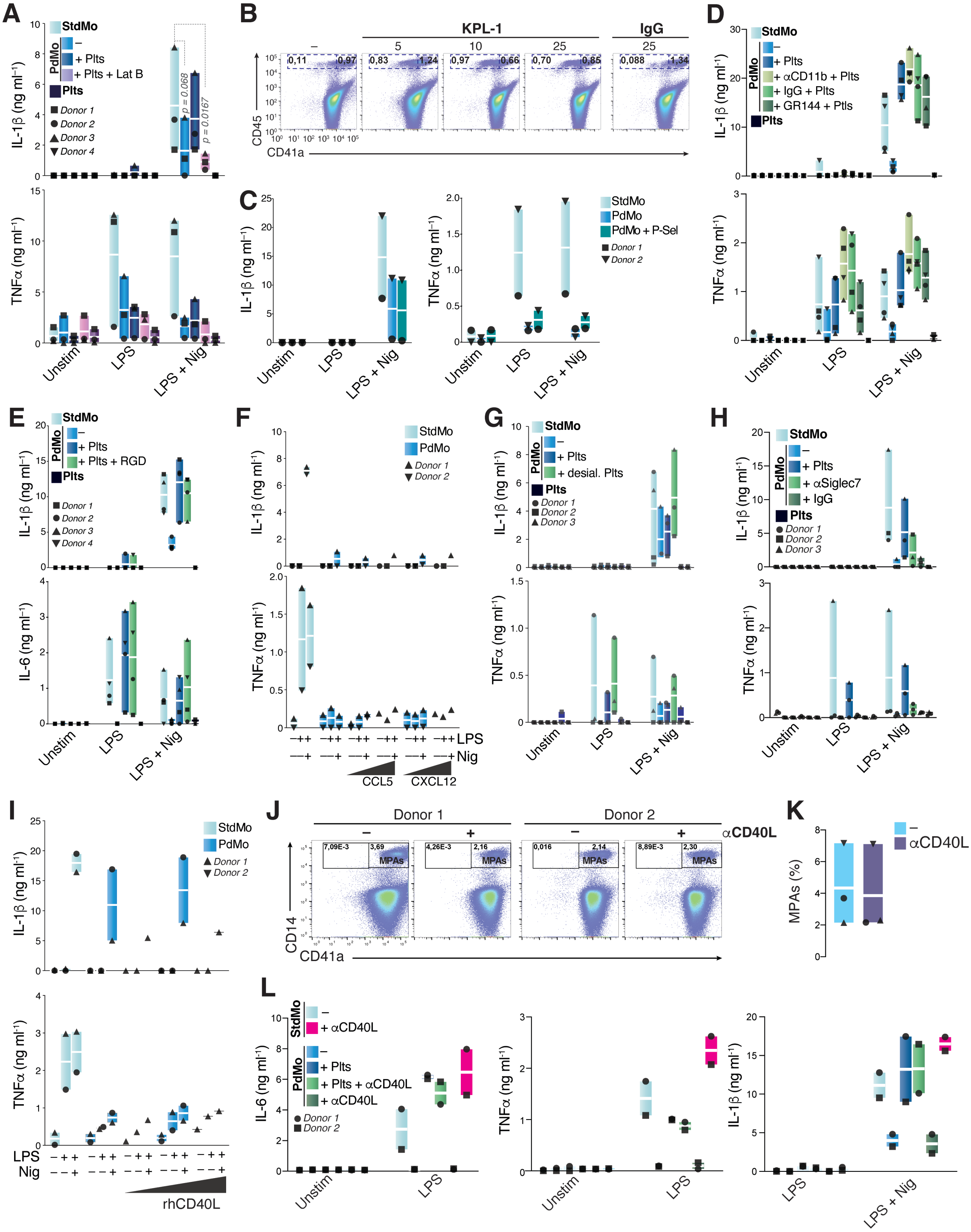
A
StdMo
PdMo
–
+ Plts
+ Plts + Lat B
Plts
Donor 1
Donor 2
Donor 3
Donor 4
IL-1β (ng ml⁻¹)
p = 0.068
p = 0.0167
TNFα (ng ml⁻¹)
Unstim
LPS
LPS + Nig
B
KPL-1
IgG
–
5
10
25
25
0,11
0,97
0,83
1,24
0,97
0,66
0,70
0,85
0,088
1,34
CD45
CD41a
C
IL-1β (ng ml⁻¹)
TNFα (ng ml⁻¹)
StdMo
PdMo
PdMo + P-Sel
Donor 1
Donor 2
Unstim
LPS
LPS + Nig
D
StdMo
PdMo
–
+ Plts
+ αCD11b + Plts
+ IgG + Plts
+ GR144 + Ptls
Plts
IL-1β (ng ml⁻¹)
TNFα (ng ml⁻¹)
Unstim
LPS
LPS + Nig
E
StdMo
PdMo
–
+ Plts
+ Plts + RGD
Plts
Donor 1
Donor 2
Donor 3
Donor 4
IL-1β (ng ml⁻¹)
IL-6 (ng ml⁻¹)
Unstim
LPS
LPS + Nig
F
StdMo
PdMo
Donor 1
Donor 2
IL-1β (ng ml⁻¹)
TNFα (ng ml⁻¹)
LPS
Nig
CCL5
CXCL12
G
StdMo
PdMo
–
+ Plts
+ desial. Plts
Plts
Donor 1
Donor 2
Donor 3
IL-1β (ng ml⁻¹)
TNFα (ng ml⁻¹)
Unstim
LPS
LPS + Nig
H
StdMo
PdMo
–
+ Plts
+ αSiglec7
+ IgG
Plts
Donor 1
Donor 2
Donor 3
IL-1β (ng ml⁻¹)
TNFα (ng ml⁻¹)
Unstim
LPS
LPS + Nig
I
StdMo
PdMo
Donor 1
Donor 2
IL-1β (ng ml⁻¹)
TNFα (ng ml⁻¹)
LPS
Nig
rhCD40L
J
Donor 1
Donor 2
αCD40L
7,09E-3
3,69
4,26E-3
2,16
0,016
2,14
8,89E-3
2,30
MPAs
CD14
CD41a
K
–
αCD40L
MPAs (%)
L
StdMo
–
+ αCD40L
PdMo
–
+ Plts
+ Plts + αCD40L
+ αCD40L
Donor 1
Donor 2
IL-6 (ng ml⁻¹)
TNFα (ng ml⁻¹)
IL-1β (ng ml⁻¹)
Unstim
LPS
LPS + Nig

**Fig S5: The Platelet-Monocyte crosstalk is independent of classical immune costimulatory molecules, Integrins, and Sialic Acids.**

**(A)** IL-1β and TNFα levels in CFS of StdMo, PdMo, and PdMo that were supplemented with autologous platelets (PdMo + Plts, 100:1 platelets/monocytes). Monocytes were pre-treated with Lacuntrulin B (Lat B, 2 μM), before being supplied with Plts. Cells were stimulated with LPS (2 ng ml-1), followed by activation with nigericin (10 μM). N = 3 biological replicates.

(**B**) Representative flow cytometry assessment and gating strategy of StdMo that were incubated with growing concentrations of KPL-1 (5, 10 or 25 μg $ml^{-1}$), or IgG (25 μg $ml^{-1}$). Gates indicate the frequencies of free platelets ($CD45^{-}$ $CD41a^{+}$), MPAs ($CD45^{+}$ $CD41a^{+}$) and platelet-free monocytes ($CD45^{+}$ $CD41a^{-}$). Data is representative of two independent experiments with several donors.

(**C**) IL-1β and TNFα levels in CFS of StdMo, PdMo, and PdMo that were supplemented with recombinant human CD62P (P-Sel, 40 ng $ml^{-1}$). Cells were stimulated with LPS (2 ng $ml^{-1}$), followed by activation with nigericin (10 μM). N = 2 biological replicates.

(**D**) IL-1β, and TNFα levels in CFS of StdMo, PdMo, PdMo, or PdMo + Plts and stimulated with LPS or LPS + Nig. PdMo were re-added with platelets in the presence of a blocking mAb against CD11b (10 μg·$ml^{-1}$), or 10μM of the inhibitor of GPIIb/IIIa (GR 144053). N = 4 biological replicates.

(**E**) IL-1β and IL-6 levels in CFS of StdMo, PdMo, PdMo, or PdMo that were supplemented with untreated (+ Plts) or platelets treated with Arginine-Glycine-Aspartate (RGD) peptides, and stimulated as in **A**. N = 4 biological replicates.

(**F**) IL-1β and TNFα levels in CFS of StdMo, PdMo, PdMo, or PdMo that were supplemented with platelets (+ Plts) or rhCXCL12 (2 - 4 ng $ml^{-1}$), or rhCCL5 (300 - 600 μg $ml^{-1}$), and stimulated as in **A**. N = 2 biological replicates.

(**G**) L-1β and TNFα levels in CFS of StdMo, PdMo, PdMo, or PdMo that were supplemented with intact (+ Plts), or desialylated platelets (desial. Ptls), and stimulated as in **A**. N = 3 biological replicates.

(**H**) IL-1β and TNFα levels in CFS of StdMo, PdMo, PdMo, or PdMo that were supplemented with untreated platelets or platelet-bound to anti-siglec7 (10 μg·$ml^{-1}$), and stimulated as in **A**. N = 3 biological replicates.

(**I**) IL-1β, TNFα, and IL-6 levels in CFS of StdMo, PdMo, or PdMo that were supplemented with viable (+ Plts), or 2% PFA-fixed platelets (+ Fixed Plts), and stimulated as in **A**. N = 4 biological replicates.

**(J)** IL-1β, and TNFα levels in CFS of StdMo, PdMo, PdMo, or PdMo + Plts and stimulated with LPS or LPS + Nig. PdMo were supplemented with platelets (+ Plts) or rhCD40L (10, 50, or 100 ng ml-1). N = 2 biological replicates.

(**K - L**) Representative flow cytometry assessment with gating strategy, and quantification (**C**) of StdMo that were incubated with a monoclonal Ab against CD40L. Gates indicate the frequencies of platelet-free monocytes ($CD45^{+}$ $CD41a^{-}$) and MPAs ($CD14^{+}$ $CD41a^{+}$). N = 2 biological replicates.

(**L**) IL-6, TNFα and IL-1β levels in CFS of StdMo, PdMo, PdMo, or PdMo that were supplemented with untreated (+ Plts) or platelets treated with 50μM of Arginine-Glycine-Aspartate (RGD) peptides, and stimulated as in **A**. As control, anti-CD40L mAb were added directly to StdMos. N = 2 biological replicates.

All graphs show floating bars display max/min values with indication to the mean (white bands). Each symbol represents one donor, or independent experiment. P values were calculated with 2-Way Anova, Tukey's multiple comparison test, and are displayed in the figures.

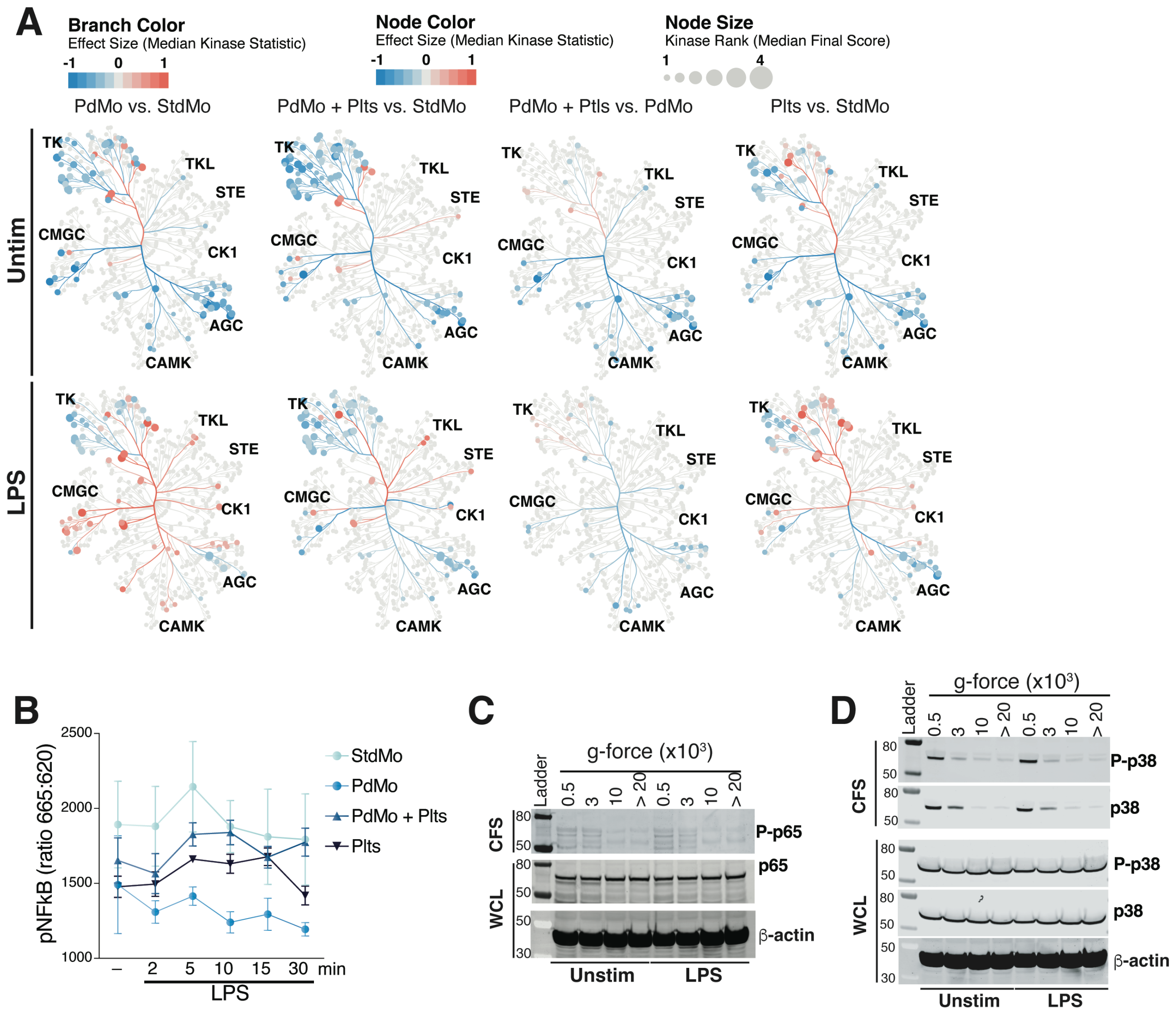


**Fig S6: Intrinsic kinase activity in platelets and their effects on human monocytes**

**(A)** Coral trees displaying the activity of Protein Tyrosine and Serine/Threonine kinases in unstimulated (Unstim) or LPS-treated (LPS) primary human monocytes comparing the effects of platelet-depletion/supplementation, measured with a PamStation12 (PamGene).

**(B)** NFκB activity assay in CFS of unstimulated (Unstim), or LPS-stimulated (100 ng ml-1, for the indicated times) primary human monocytes (StdMo), PdMo, PdMo + Plts, and Plts alone. Cells were lysed/incubated with lysis buffer, supplied Cisbio, and used to assess phosphorylated NFkB by HTRF (Cisbio). Graphs show floating bars display max/min values with indication to the mean (white bands). Each symbol represents one donor, or independent experiment.

**(C - D)** Immunoblot of RelA (p65 and p-p65) (**C**) or (p-38 and P-p38) (**D**) in resting (Unstim) or LPS-activated (LPS) human platelets. Platelets were submitted to centrifugation at 500, 3000, 10,000, or 20,000 x g and the levels of proteins were assessed in the pellets (WCL) or supernatants (CSF) after centrifugation. Results are representative of two independent experiments.

### Unstim Monocytes + SILAC MEG-01 Sups (n = 6)

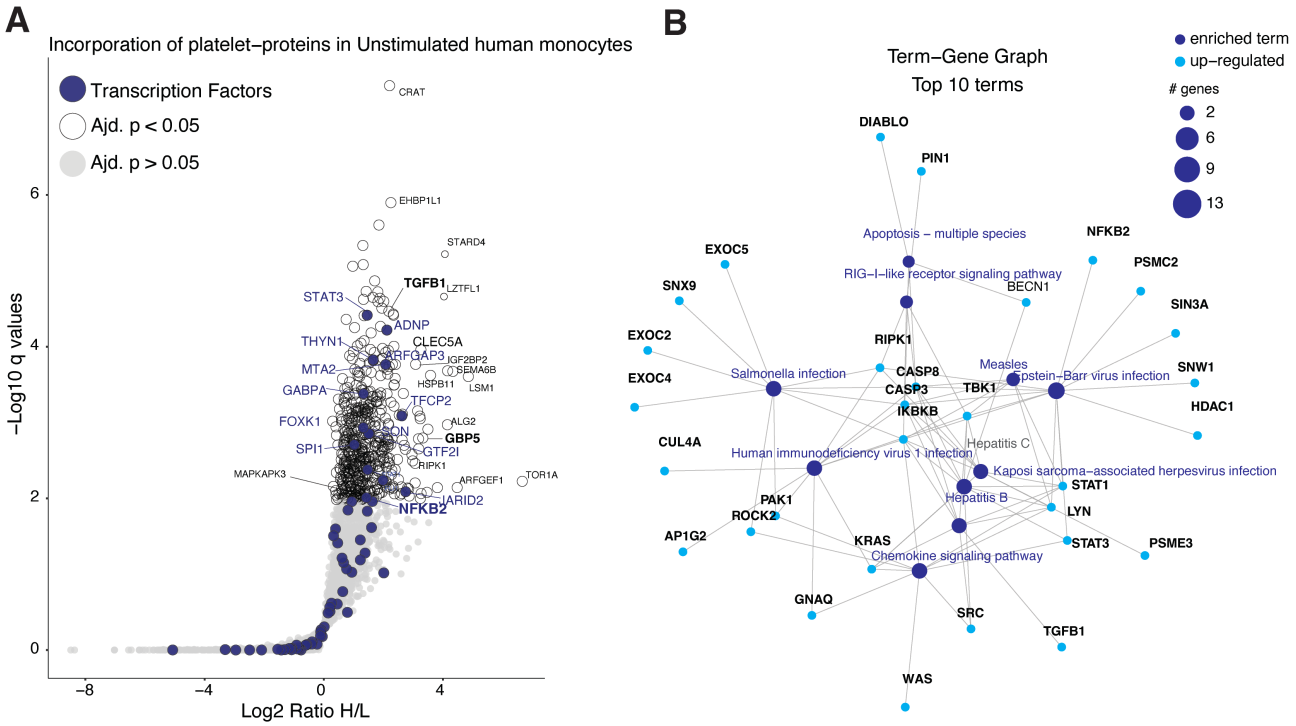


**Fig S7: Analysis of Mass speck proteomics combined with Stable isotope labeling with amino acids in cell culture (SILAC)**

**(A)** Volcano plots of proteins with heavy-AAs detected in PdMo exposed to cell-free supernatants of MEG-01 cells through Mass speck proteomics combined with Stable isotope labeling with amino acids in cell culture (SILAC). Unstimulated conditions are shown (n = 6 biological replicates). LPS-stimulated conditions are shown in Fig 6.

**(B)** Pathway analysis of the proteins with heavy-AAs detected in PdMo exposed to cell-free supernatants of MEG-01 cells as in A.

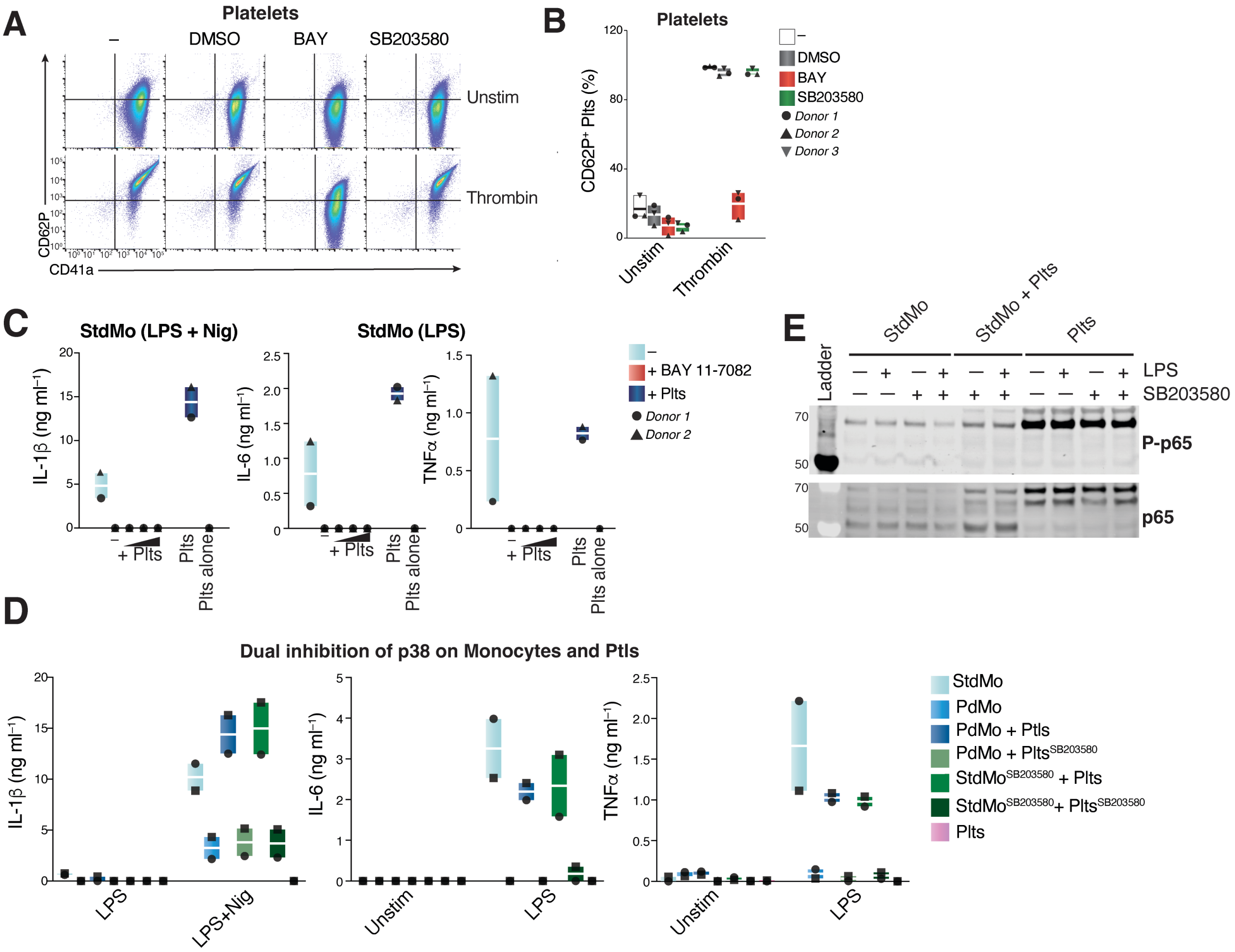


**Supplemental Fig S8: Inhibition of p38 MAPK and NFkB in platelets, human monocytes, or both.**

**(A - B)** Representative flow cytometry assessment and (**B**) quantification (%, n = 3) of CD41a and P-selectin (CD62P) expression on human platelets that were pre-treated with BAY, or SB203580 before being activated with Thrombin (1 U ml⁻¹).

(**C**) IL-1β, TNFα, and IL-6 concentrations in the CFS of StdMo that were treated with BAY before been added with increasing ratios of freshly isolated platelets (1:5, 1:50, and 1:100). Cells were stimulated with LPS or LPS and nigericin (LPS + Nig). Floating bars display max/min values with indication to the mean (white bands). Each symbol represents one donor.

**(D)** Concentrations of IL-1β, IL-6, and TNFα in CFS of StdMo, PdMo, or PdMo that were supplemented with platelets pretreated with SB203580 (Plts + SB), or left untreated (+ Plts), or in StdMo pre-treated with SB203580 (StdMo + SB) that were supplemented with intact platelets (StdMo + SB + Plts) or with SB203580-treated platelets (StdMo + SB + Plts + SB). Co-cultures were stimulated with LPS (2 ng/ml) followed by nigericin stimulation (10 µM). Floating bars display max/min values with indication to the mean (white bands). Each symbol represents one donor, (n = 2).

(E) Immunoblotting for phospho-p65 and total p65 (RelA) on unstimulated and LPS-stimulated StdMo primary human monocytes treated with 50 µM of SB203580.
